# T-bet Expressing B cells are Key Determinants of Protective Immunity Against Norovirus Infection

**DOI:** 10.64898/2026.03.30.715451

**Authors:** Abeera Mehmood, Taylor Basso, Alana Weiss, Stanislav Dikiy, Kyong T. Fam, Daniella Marinelli, Susanna Manenti, Nicole Wolman, Meng Zhang, Bryan Briney, Howard C. Hang, Alejandra Mendoza

## Abstract

The gastrointestinal tract (GI) is the largest environmental mucosal interface and is exposed to diverse commensal and pathogenic microbes. B cells are a prevalent immune component of the GI tract and its associated secondary lymphoid organs, yet we know little about the diversity and stability of distinct transcriptional programs that modulate B cell responses against different classes of pathogens or environmental perturbations. A subset of B cells defined by expression of the transcription factor T-bet, has been canonically associated with antiviral immunity through IgG production. However, the role of T-bet expressing B cells in mucosal tissues, where IgA responses predominate, is poorly understood. Here, we identify a population of intestinal T-bet+ B cells that, in the absence of overt perturbation, constitutes a minor fraction of intestinal associated B cells and undergoes continuous turnover. In contrast, during enteric viral infection with murine norovirus (MNV), T-bet⁺ B cells undergo a marked expansion, with T-bet expression stably maintained in the majority of virus-specific B cells, including IgG2c and IgA switched B cells. Moreover, virus-reactive IgG2c and IgA B cells arise independently rather than through sequential switching, and B cell intrinsic T-bet expression is required for effective germinal center responses but dispensable for IgA class switching. Moreover, T-bet expressing B cells are required for the generation of all MNV-specific circulating IgG and mucosal IgA, and for protection upon re-encounter with the virus. Together, these findings establish T-bet expressing B cells as a specialized B cell subset essential for mucosal immunity and protection against norovirus infection.

## INTRODUCTION

The gastro-intestinal (GI) tract is a complex and unique tissue exposed to a diverse array of stimuli that drive the activation, differentiation, and functional specialization of immune cells^1,2^. Among immune populations in the GI tract, B cells are among the most prevalent and are responsible for antibody production, a defining feature of vertebrate adaptive immunity that enables durable, secreted, and finely tuned responses against pathogens^3,4^. While we understand many facets of B cell biology, many questions about their functions remain unanswered, especially those pertaining to their adaptations to and residence in mucosal tissues where high rates of antibody production take place^3,5^.

While antibody isotypes have distinct effector functions, even antibodies of the same isotype can perform diverse and sometimes non-overlapping functions^6,7^. For instance, IgA dominates at mucosal surfaces and plays key roles in both limiting pathogen invasion and maintaining host-microbiota homeostasis^7,8^. IgG can also contribute to protection against enteric pathogens and help restrict systemic spread of infection^9,10^. Consistently, germinal centers in the murine intestine contain B cell populations that have switched to mixed isotypes, suggesting that mucosal immune responses can give rise to multiple antibody classes concurrently or sequentially^11–15^. These observations suggest that effective mucosal immunity depends on coordinated production of multiple antibody classes.

In addition, beyond class-switching, B cell function can also be shaped by specialized programs that modulate functions like tissue residency, longevity, and antibody affinity^16–19^. Given that the intestine is a major site of entry and a reservoir for many viral pathogens^20^, specialized transcriptional B cell programs are likely critical for mounting effective protective immunity at this barrier tissue. A specialized transcriptional program in B cells is defined by expression of the transcription factor T-bet, encoded by *Tbx21*, a key regulator of type 1 immune responses^21^. B cells expressing T-bet are induced in response to viral infection, and these cells are required for antiviral IgG production^22–25^. Moreover, T-bet itself is required in B cells for isotype switching to IgG2c/a (in mice)^26^ and IgG1 and IgG3 (in humans)^25^. Beyond infection, T-bet expressing B cells have also been reported to expand during aging and autoimmunity, suggesting that this population is also responsive to additional uncharacterized immunological stimuli^27–31^. Yet, a role for this subset of B cells in the GI mucosa has not been reported to date. Here, we show that T-bet-expressing B cells comprise the dominant virus-specific B cell population during enteric viral infection. These cells generate the majority of antigen-specific germinal center and memory B cell responses and are required for the production of antiviral antibodies in circulation and at mucosal sites. Genetic ablation of T-bet-expressing B cells results in a profound loss of virus-specific IgG and IgA responses and compromises protective immunity upon subsequent viral exposure. Together, these findings identify T-bet⁺ B cells as a durable antiviral immune cell population that persists after infection and coordinates systemic and mucosal antibody immunity against enteric viruses.

## RESULTS

### Intestinal T-bet⁺ B cells continuously turn over in response to local tissue cues

Given the constant and diverse antigenic exposure within the intestine and the marked regional variation in the local intestinal environment^2^, we asked whether T-bet is expressed in B cells throughout the GI tract at homeostasis. Using knock-in mice carrying IRES-tdTomato-T2A-Cre in the 3′ UTR of *Tbx21 (Tbx21^tdTomato-cre^)*^32^, we used tdTomato to examine T-bet expression in B cells along the GI tract. We found tdTomato expression across all B cell subsets including germinal center (GC) B cells, memory B cells (MBC), follicular B cells (FoB), and Plasma cells **(Fig. 1a-b, Extended data Fig. 1a-c)**. Within the intestinal lamina propria, tdTomato was expressed in a small fraction of plasma cells (less than 5%), while a larger proportion of MBC express tdTomato, increasing in frequency towards the distal small intestine and colon **(Fig. 1a)**. As Peyer’s Patches (PP) can directly sample antigens through M cells, they are specialized for localized responses to antigens sampled from the intestinal lumen^33^. Therefore, the cellular composition of PP along the length of the intestine reflects regional cues that shape immune programs. Consistent with distally increasing T-bet expressing B cells in the lamina propria, mesenteric lymph nodes (mLN) and distal PP also harbored higher frequencies of tdTomato expressing GC B cells and MBC, while the proportion tdTomato+ FoB was relatively low throughout the length **(Fig. 1b)**. Although T-bet is classically linked to IgG2c class switching^26^, we found that in the absence of infection, a fraction of IgA⁺ MBC and GC B cells within PP and mLN express tdTomato, as did IgA⁺ MBC and plasma cells in the lamina propria **(Fig. 1c-d)**. These observations indicate that T-bet expression marks a subset of B cells across the GI tract and its associated secondary lymphoid organs, including IgA switched cells. Moreover, the proportion of T-bet⁺ B cells increased progressively along the GI tract, suggesting that local environmental cues may drive T-bet induction in B cells or sustain their persistence within these tissues.

**Figure 1.**
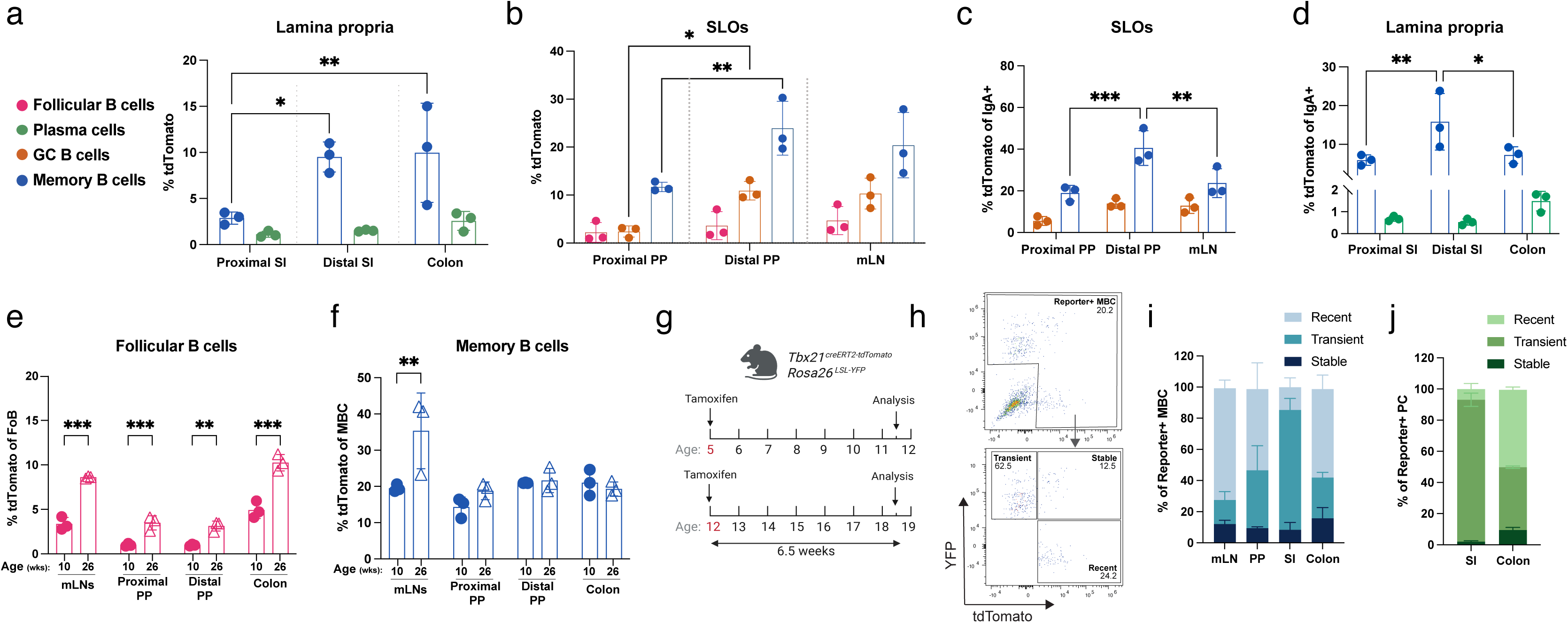
Intestinal T-bet⁺ B cells continually turn over in response to local tissue cues. **(a-f)** Secondary lymphoid organs (SLOs) and lamina propria from uninfected *Tbx21^tdTomato-cre^* mice were analyzed by flow cytometry. (**a**) Frequency of tdTomato+ memory B cells (MBC) and plasma cells in the proximal small intestine (SI), distal SI, and colon lamina propria. (**b**) Frequency of tdTomato+ GC B cells, MBC, and Follicular B cells in proximal PP, distal PP, and mLN. (**c**) Frequency of tdTomato+ IgA-switched GC B cells and memory B cells in PP and mLN. (**d**) Frequency of tdTomato+ IgA-switched memory B cells and plasma cells in proximal small intestine (SI), distal small intestine (SI), and colon lamina propria. **(a-d)** Mice were analyzed at 8 weeks of age. (**e**) Frequency of tdTomato+ Follicular B cells (FoB) in mLN, PP, and colon lamina propria of 10- and 26-week-old mice. (**f**) Frequency of tdTomato+ memory B cells (MBC) in mLN, PP, and colon lamina propria of 10- and 26-week-old mice. **(g-j)** Uninfected *Tbx21^tdTomato-creERT2^ Rosa26^LSL-YFP^* were administered tamoxifen by oral gavage at 5 or 12 weeks of age and analyzed by flow cytometry 6.5 weeks following tamoxifen administration. (**g**) Schematic for assessing stability of T-bet expression in *Tbx21^tdTomato-creERT2^ Rosa26^LSL-YFP^* mice used for h-i. (**h**) Representative flow cytometry plots pre-gated on memory B cells (MBC). Flow plots show gating strategy for identifying T-bet recent, T-bet transient and T-bet stable B cells within all reporter+ cells (YFP+ and tdTomato+) (**i**) Frequency of T-bet Recent (tdTomato+ YFP-), T-bet transient (tdTomato- YFP+), and T-bet stable (tdTomato+ YFP+) cells within reporter+ memory B cells (MBC) from mice given a tamoxifen pulse at 12 weeks of age. (**j**) Frequency of T-bet Recent (tdTomato+ YFP-), T-bet transient (tdTomato- YFP+), and T-bet stable (tdTomato+ YFP+) cells within reporter+ plasma cells (PC) from mice given a tamoxifen pulse at 12 weeks of age. **(a-f)** Each dot represents a mouse, bars show mean, error bars show S.D; 2-way ANOVA with Tukey’s correction for multiple comparisons. **(i-j)** Bars show mean, error bars show S.D. n=3. *P< 0.033(*), P< 0.002(**), P < .001(***)*.

Several studies in both mice and humans have reported age-associated increases in T-bet-expressing B cells^27–29^. To assess whether aging alters mucosal B cell compartments, we profiled tdTomato⁺ B cells in mice of different ages. Older mice (26 weeks old) exhibited modest increases in the frequency of tdTomato⁺ FoB cells across intestinal sites and an increase in tdTomato⁺ MBC specifically within mLN relative to younger mice (10 weeks old) **(Fig. 1e-f)**. Importantly, this expansion was not observed uniformly across all B cell subsets or across all intestinal sites **(Extended data Fig. 1d-e)**, indicating that aging, in the absence of overt inflammatory perturbation, induces a limited and subset-restricted accumulation of T-bet⁺ B cells in the GI tract and associated secondary lymphoid organs.

To determine whether T-bet⁺ B cells represent a long-lived resident population or are continuously turned-over within the intestinal environment, we fate-mapped T-bet expression in B cells in mice carrying tamoxifen inducible creERT2 and tdTomato in the 3′ UTR of *Tbx21 (Tbx21^tdTomato-creERT2^)*^32^ and a YFP recombination reporter (*Rosa26^LSL-YFP^*). Tamoxifen was administered to several cohorts of mice by oral gavage at 5 weeks or at 12 weeks of age, thereby permanently labeling all cells expressing T-bet at the time of gavage with YFP **(Fig. 1g)**. All mice were analyzed 6.5 weeks after tamoxifen labeling to assess the persistence of T-bet expression over time **(Fig. 1g)**. We analyzed intestinal and lymphoid tissues to distinguish: T-bet Transient (YFP⁺, tdTomato⁻) cells that lost tdTomato expression after the tamoxifen pulse, T-bet Stable (YFP⁺, tdTomato⁺) cells that maintain tdTomato expression, and T-bet Recent (YFP⁻ tdTomato⁺) cells that up-regulate tdTomato after the labeling window **(Fig. 1h)**. Across all cohorts, the majority of tdTomato⁺ B cells lacked YFP labeling or had lost tdTomato expression, indicating that most T-bet⁺ B cells are newly generated and those that stay in the tissue are either short lived or lose T-bet expression **(Fig. 1i-j, Extended data Fig. 1f-g)**. Indeed, only a small fraction of YFP⁺ cells retained tdTomato expression 6.5 weeks after labeling, with the highest retention observed among colonic MBC and plasma cells **(Extended data Fig. 1h-j)**. These data indicate that intestinal T-bet⁺ B cells undergo constant turnover under homeostatic conditions with only a minor fraction of cells becoming long-lived. Collectively, these results demonstrate that T-bet⁺ B cells form a small dynamic and regionally enriched population of intestinal B cells.

### T-bet expressing B cells are responsive to changes in the microbiota

As the microbiota represents a major source of stimulation in the intestine, we next asked if intestinal commensal microbes drive the induction of T-bet⁺ B cells. To deplete intestinal bacteria, *Tbx21^tdTomato-cre^*mice were treated with oral antibiotics for 2 weeks and then analyzed **(Fig. 2a)**. Antibiotic treatment differentially altered specific B cell populations across intestinal compartments, including T-bet⁺ B cells, indicating that its effects were not restricted to this subset alone. Within lymphoid organs, distal PP showed reduced GC B cells and MBC, proximal PP remained largely unchanged, whereas mLN displayed increased GC B cells with no changes in MBC **(Fig. 2b-c)**. In the lamina propria, total plasma cells, including IgA⁺ plasma cells were reduced in both the small intestine and colon, with a stronger reduction in the small intestine **(Fig. 2d-e)**. MBC within the lamina propria remained largely unchanged (**Extended data Fig. 2a)**. T-bet⁺ B cells followed the same pattern as bulk B cells. tdTomato⁺ GC B cells were reduced following antibiotic treatment, with a pronounced decrease in distal PP and a more modest trend in proximal PP **(Fig. 2f)**. tdTomato+ MBC in PP exhibited a similar pattern **(Fig. 2g)**, suggesting that the induction of T-bet in some subsets of B cells is sensitive to perturbations in the microbiota. In contrast, within mLN the number of tdTomato⁺ cells remained unchanged, while the frequency of T-bet⁺ GC B cells and MBC decreased, likely reflecting the increase in total GC B cells and a numerical increase in total MBC in this compartment **(Fig. 2f-g, Extended data Fig. 2b-c)**. Within the lamina propria, the frequency of tdTomato+ MBC and plasma cells, including IgA⁺ plasma cells, remained unchanged **(Fig. 2h-i, Extended data Fig. 2d)**. The stability of T-bet⁺ plasma cells is consistent with previous reports where adult mice show little change in the IgA repertoire even several weeks after antibiotic treatment^34^. Together, these data suggest that in the absence of infection, the microbiota contributes to the induction of T-bet expression in a small subset of intestinal B cell populations but is dispensable for the short-term maintenance of T-bet⁺ plasma cells present in the tissue.

**Figure 2.**
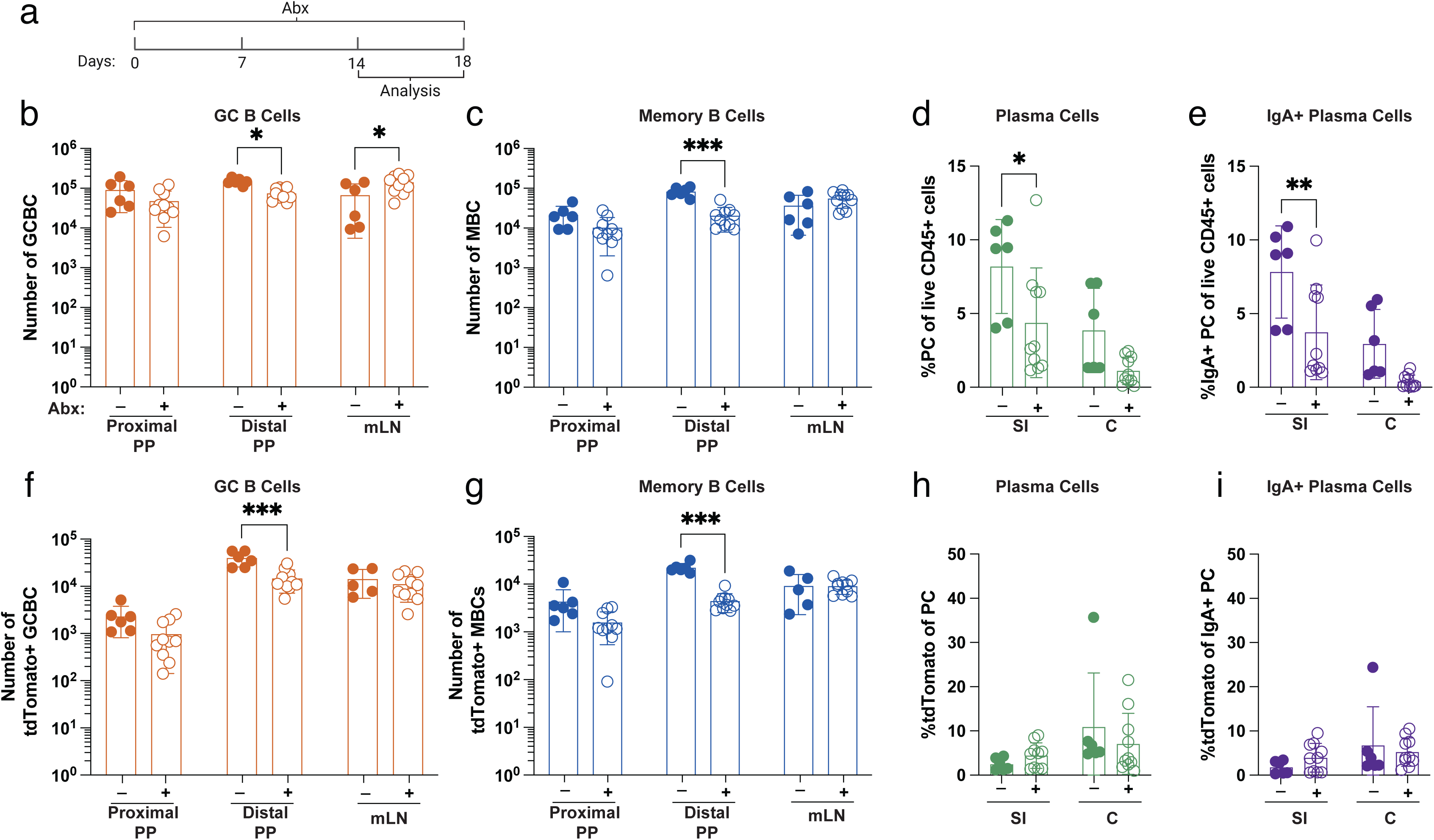
T-bet expressing B cells are responsive to changes in the microbiota. *Tbx21^tdTomato-cre^*mice were administered a triple antibiotic cocktail in drinking water for 14 days and analyzed by flow cytometry. **(a)** Schematic of antibiotic administration and time of analysis. **(b)** Number of GC B cells (GCBC) in mLN and PP. **(c)** Number of memory B cells (MBC) in mLN and PP **(d)** Frequency of plasma cells (PC) in small intestine (SI) and colon lamina propria. **(e)** Frequency of IgA⁺ plasma cells (PC) in small intestine (SI) and colon lamina propria. **(f)** Number of tdTomato+ GC B cells (GCBC) in mLN and PP. **(g)** Number of tdTomato+ memory B cells (MBC) in mLN and PP. **(h)** Frequency of tdTomato+ plasma cells (PC) in small intestine (SI) and colon lamina propria. **(i)** Frequency of tdTomato+ of IgA⁺ plasma cells (PC) in small intestine (SI) and colon lamina propria. **(b-g)** Each dot represents a mouse, bars show mean, error bars show S.D; 2-way ANOVA with Bonferroni correction for multiple comparisons *P< 0.033(*), P< 0.002(**), P < .001(***)*.

### T-bet expressing B cells are induced in response to enteric viral infection

Given that under homeostatic conditions T-bet⁺ B cells constitute only a small fraction of intestinal B cells, we sought to determine whether enteric viral infection drives their expansion within the GI tract. Among enteric viruses, noroviruses are currently the leading cause of viral gastroenteritis and pose a significant health risk to children and immune compromised individuals^35^. Murine norovirus (MNV) is a natural enteric pathogen that models key immunological features of human norovirus infection^36^. B cells play an essential role in clearing MNV infection as B cell deficient mice are unable to clear acute MNV infection^9^. Thus, we asked whether T-bet-expressing B cells contribute to the response against MNV infection. To compare B cell responses to acute infection we used an acute MNV strain (MNV-CW3), cleared within 7 days in immune competent mice^9^. In response to MNV infection of *Tbx21^tdTomato-cre^* mice, we observed increased frequencies of tdTomato expressing B cells in mLN and PP at 14 days post infection **(Fig. 3a-b)**. The increase in tdTomato expression was specific to activated B cell subsets, including GC B cells and MBC, and was not observed in FoB **(Fig. 3a-b)**. Consistent with these findings, imaging of mLN revealed enlarged germinal centers (IgD- B220+) populated with tdTomato+ GC B cells in the mLN of infected mice compared with uninfected (PBS) controls **(Fig. 3c)**.

**Figure 3.**
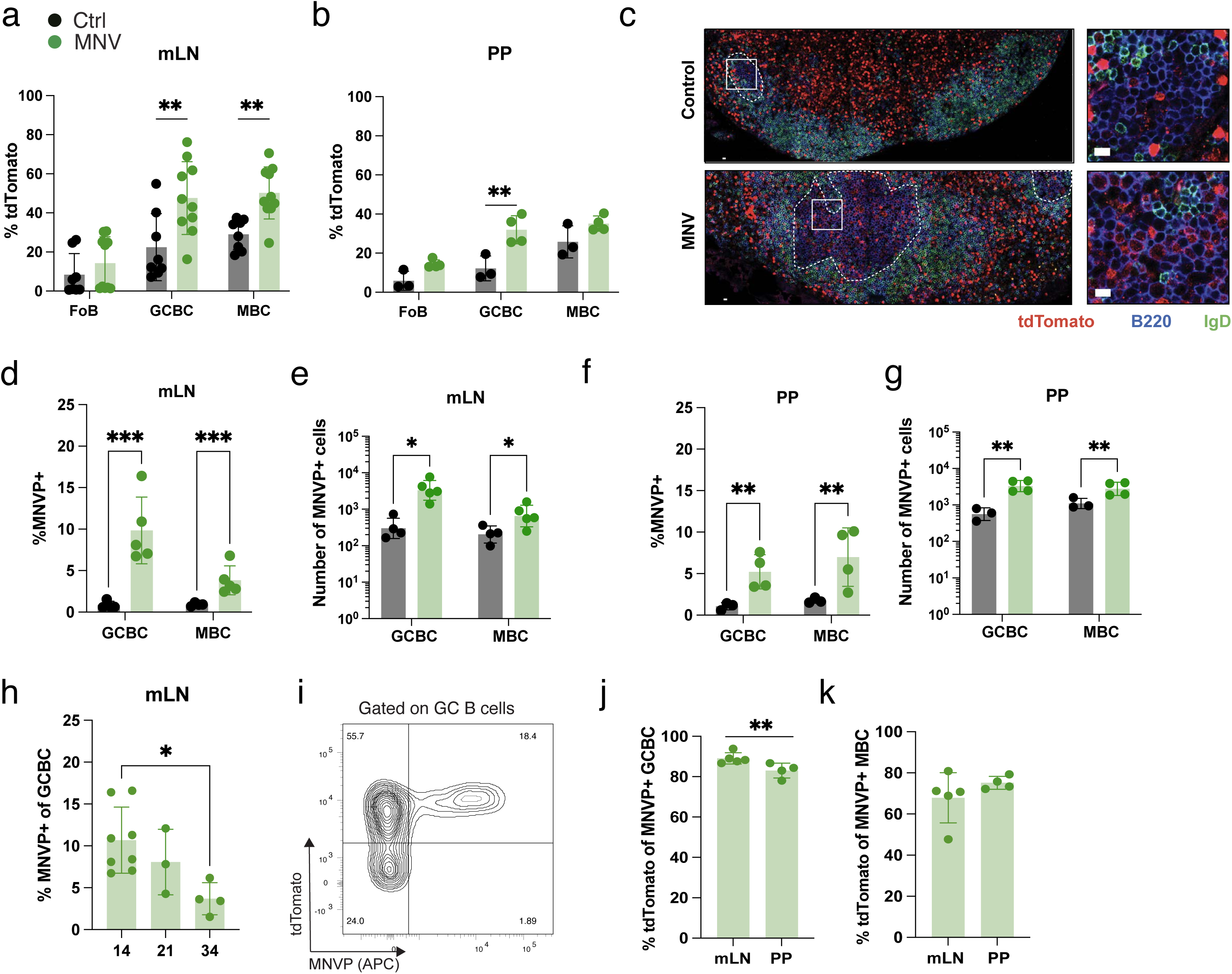
T-bet expressing B cells are induced in response to enteric viral infection. **(a-c)** *Tbx21^tdTomato-cre^* mice were infected with 1E6 PFU of MNV and lymphoid organs profiled at 14 dpi by flow cytometry and microscopy. **(a)** Frequency of tdTomato+ B cell subsets in mLN. **(b)** Frequency of tdTomato+ B cell subsets in PP. **(c)** Representative confocal micrograph of infected and control mLN. Germinal center B cells were defined as B220+IgD-. Germinal centers are outlined by dashed line. Box shows are of inset. Scale bars, 10 μm. tdTomato (red), B220 (blue), IgD (green). **(d-k)** *Tbx21^tdTomato-cre^* mice were infected with 1E6 PFU of MNV and lymphoid organs profiled for reactivity to MNVP probe and tdTomato expression by flow cytometry at 14 dpi or at indicated timepoints. **(d)** Frequency of MNVP+ GC B cells (GCBC) and memory B cells (MBC) in mLN at 14 dpi. **(e)** Number of MNVP+ GC B cells and MBC in mLN at 14 dpi. **(f)** Frequency of MNVP+ GC B cells and MBC in PP. **(g)** Number of MNVP+ GC B cells and MBC in PP. **(h)** Frequency of MNVP+ of GC B cells in mLN over indicated time points post infection. **(i)** Representative flow cytometry plot showing MNVP vs tdTomato on MNVP+ GC B cells in mLN at 14 dpi**. (j)** Frequency of tdTomato+ MNVP+ GC B cells and **(k)** MBC. **(a-b, d-h, and j-k)** Each dot represents a mouse, bars show mean, error bars show S.D; **(a-b, d-g)** 2-way ANOVA with Bonferroni correction for multiple comparisons; **(h)** One-way ANOVA with Bonferroni correction for multiple comparisons; **(j-k)** Unpaired t-tests. *P< 0.033(*), P< 0.002(**), P < .001(***)*.

To assess whether T-bet is induced in antigen specific B cells, we recombinantly expressed and purified the P domain of the surface exposed VP1 capsid protein from the MNV-CW3 strain, hereafter referred to as MNVP. The P domain is considered the most variable region of the capsid and serves as the major site for host factor engagement and antibody binding^37–42^. Residues 229-541 of VP1 were C-terminally fused to a HisTag and AviTag to enable purification and biotinylation, respectively **(Extended data Fig. 3a).** Recombinant MNVP was efficiently biotinylated enabling subsequent tetramerization using fluorescently conjugated streptavidin to generate a probe for identifying MNVP-specific B cells **(Extended data Fig. 3b-c).** The resulting fluorescent MNVP probe efficiently labeled MNVP-specific B cells within mLN and PP by flow cytometry and enabled detection of MNVP-specific antibodies in serum, with minimal background staining in uninfected mice **(Fig 3d-g, Extended data Fig. 3d).** To validate the specificity of MNVP flow staining, we performed dual-color tetramer staining using MNVP complexed with two distinct fluorophores and confirmed that antigen-specific B cells were positive for both fluorophores **(Extended data Fig. 3e).** We detected MNVP-specific GC B cells and MBC at comparable numbers in the PP and mLN 14 days post-infection, indicating that B cell priming by MNV infection occurs at both sites **(Fig. 3e & 3g**). Reflecting the overall higher abundance of GC and MBC present in PP compared to mLN, we found that approximately 10% of GC B cells and 4% of MBC in the mLN were MNVP-specific while, within PP 5% of GC B cells and 7% of MBCs were MNVP+ **(Fig. 3d & 3f**). These data indicate that the MNVP-specific germinal center response and MBC can be found in the mLN and PP. In addition, we observed that the MNVP-specific GC response in the mLN peaked at 14 days post infection and was maintained for at least 34 days **(Fig. 3h).** Next, we sought to analyze T-bet expression on MNVP-specific B cells and found that the majority of MNVP-specific GC B cells and MBC in the mLN and PP express tdTomato with approximately 85% of GC B cells and 75% of MBC expressing tdTomato **(Fig. i-k**). Together, these data demonstrate that enteric viral infection robustly induces T-bet expression in antigen-specific B cells, linking enteric antiviral immune responses to the induction of a distinct transcriptional program in intestinal B cell populations.

### Independently arising IgA⁺ and IgG2c⁺ MNV-specific B cells both express T-bet

During MNV infection, antibodies are able to confer protection as exogenous administration of MNV capsid specific neutralizing IgG antibodies reduces viral titers in infected *Rag1*^-/-^ mice^9^. As the primary site of MNV infection is the intestinal mucosa, IgA is likely also critical during MNV infection. Consistent with this, IgA is important for protection against rotavirus, and human norovirus studies have linked IgA responses with reduced viral loads and decreased disease severity, suggesting that IgA may similarly help control norovirus infection^8,43,44^. Indeed, we find that in response to MNV infection, there is robust induction of serum IgG, whereas within the intestinal lumen the MNV-specific antibody response is IgA-dominant, with little to no detectable IgG **(Fig. 4a-b).** Consistent with a known role for T-bet in IgG2c class switching^19,22,24,45,46^, we found tdTomato expression in IgG2c-switched GC B cells and MBC during MNV infection in both the mLN and PP **(Fig. 4c-d).** In addition, we also found tdTomato expression in IgA-switched GC B cells and MBC in both mLN and PP in response to MNV infection **(Fig. 4e-f).** Within the antigen specific B cell compartment, more than 50% of the MNVP-specific GC B cells and MBC in the mLN are switched to IgG2c with IgA making up only 5-10% of the response **(Fig. 4g-h).** While in the PP, IgA dominates the MNVP-specific GC B cells and MBC response followed by 10% of the response being IgG2c **(Fig. 4g-h).** This is consistent with other enteric infections like rotavirus which also induces a stronger IgA response in the PP^47^. Of note, MNVP-specific B cells also included B cells switched to IgG1 and IgG2b, albeit at lower frequencies **(Extended data Fig. 4a).** Importantly, we found that the majority of MNVP-specific IgA and IgG2c switched GC B cells and MBC express T-bet as we found that these populations were positive for tdTomato **(Fig. 4i-k)**.

**Figure 4.**
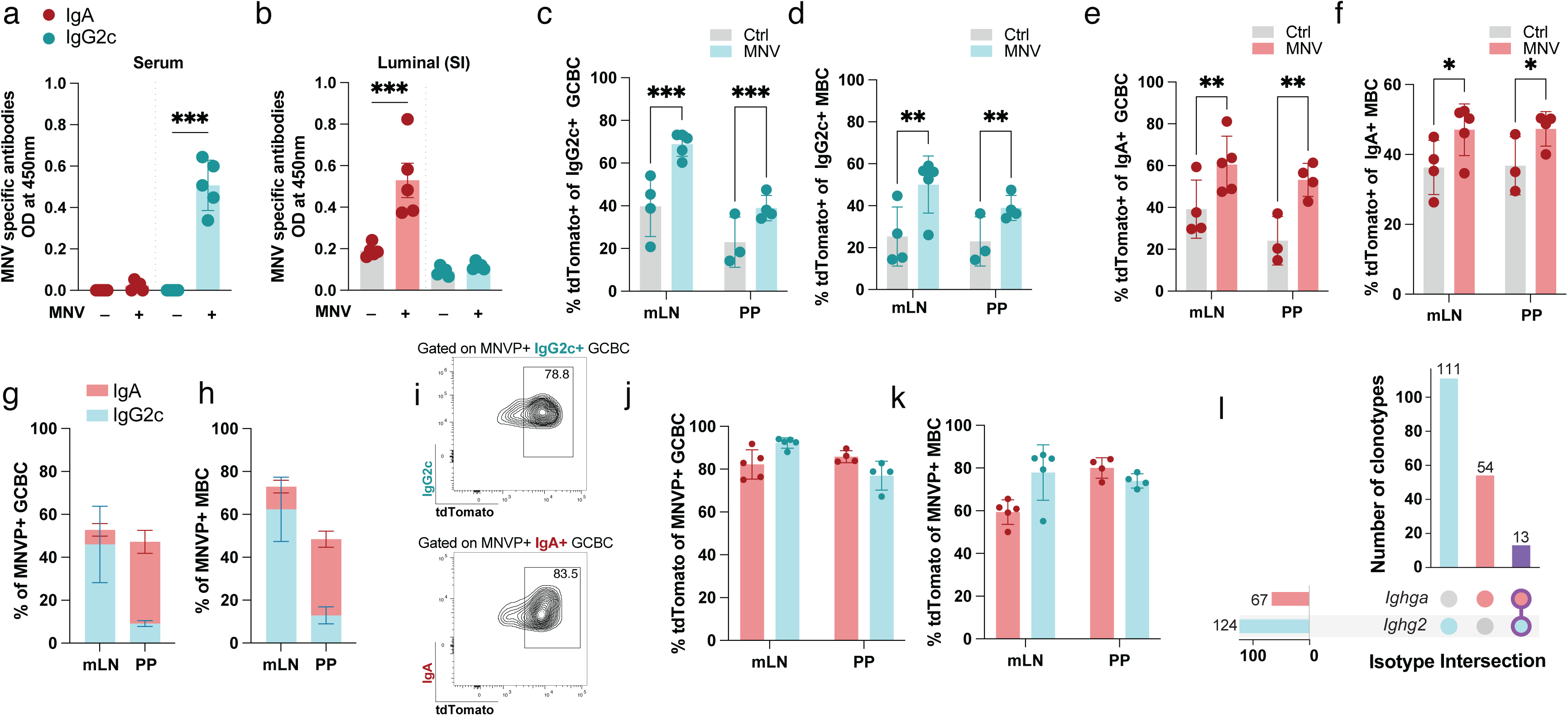
Independently arising IgA⁺ and IgG2c⁺ MNV-specific B cells both express T-bet. **(a-b)** Mice were infected with 1E6 PFU of MNV, MNV-specific IgA and IgG titers in **(a)** serum and **(b)** small intestine (SI) luminal contents 21 dpi were analyzed by ELISA. **(c-j)** *Tbx21^tdTomato-cre^* mice were infected with 1E6 PFU of MNV and lymphoid organs were analyzed by flow cytometry at 14 dpi. **(c)** Frequency of tdTomato+ IgG2c⁺ GC B cells (GCBC) from mLN and PP. **(d)** Frequency of tdTomato+ IgG2c⁺ memory B cells (MBC) in mLN and PP. **(e)** Frequency of tdTomato+ IgA⁺ GC B cells (GCBC) in mLN and PP. **(f)** Frequency of tdTomato+ IgA⁺ memory B cells (MBC) in mLN and PP. **(g)** Frequency of IgA⁺ and IgG2c⁺ MNVP+ GC B cells (GCBC) in mLN and PP. **(h)** Frequency of IgA⁺ and IgG2c⁺ MNVP+ memory B cells (MBC) in mLN and PP. **(i)** Two dimensional flow cytometry plot showing tdTomato expression on MNVP+ IgG2c and IgA-switched GC B cells (GCBC) from PP. **(j)** Frequency of tdTomato+ IgA⁺ and IgG2c⁺ MNVP+ GC B cells (GCBC) in mLN and PP. **(k)** Frequency of tdTomato+ IgA⁺ and IgG2c⁺ MNVP + MBC in mLN and PP 14 dpi. **(l)** Mice were infected with 1E6 PFU of MNV, and IgA⁺ and IgG2c⁺ MNVP⁺ GC and memory B cells were sorted from the mLNs of 6 mice at 14 dpi, pooled, and analyzed for B cell receptor sequences by scRNA-seq. Isotype intersection plots show shared clonotypes between MNVP⁺ IgG2c switched, and IgA switched B cells. **(a-f & j-k)** Each dot represents a mouse, bars show mean, error bars show S.D; 2-way ANOVA with Bonferroni correction for multiple comparisons. **(g-h)** Bars represent mean, error bars show S.D, n=4. *P< 0.033(*), P< 0.002(**), P < .001(***)*.

Due to the linear organization of the immunoglobulin heavy chain locus, class-switch recombination can occur sequentially, whereby B cells undergo successive switching events through intermediate isotypes^48,49^. Downstream isotypes in the locus, the last being IgA, can therefore arise from B cells that have previously switched to upstream isotypes^11,49^. Prior studies using oral immunization, where IgG1⁺ cells dominate the antigen-specific response, have demonstrated sequential switching from IgG1 to IgA^12^. As T-bet is induced on both IgG2c⁺ and IgA⁺ MNV-specific B cells, and IgG1⁺ B cells constitute only a minor fraction of the MNVP-specific response (**Fig. 4i-j, Extended data Fig. 4a**), we next asked whether IgA⁺ cells might instead arise from previously IgG2c-switched cells through sequential class switching. To address this, we performed B cell receptor (BCR) sequencing (BCR-seq) to assess clonal relationships between IgG2c⁺ and IgA⁺ MNVP-specific B cells. We observed minimal clonal overlap between these populations, with 86% of IgG2c⁺ and IgA⁺ clones showing no shared lineage **(Fig. 4l)**. Together, these data indicate that during enteric viral infection, T-bet is induced on both IgG2c⁺ and IgA⁺ B cells, but these populations arise largely independently. Given that class-switch recombination can occur prior to germinal center entry, our findings suggest that IgG2c⁺ and IgA⁺ B cells may be specified early through distinct differentiation pathways rather than through sequential switching^50^. In this context, T-bet expression in both populations likely reflects a shared activation or environmental cue rather than a direct role in directing a common lineage trajectory between these isotypes.

### Enteric viral infection drives stable B cell T-bet expression

Given that T-bet expression in intestinal B cells is highly dynamic at homeostasis, with rapid turnover and continual induction in newly generated cells, we next asked whether T-bet expression induced during enteric viral infection is similarly transient or instead stably maintained. To do so, we fate-mapped T-bet expression for up to 12 weeks following MNV infection in *Tbx21^tdTomato-creERT2^* mice also carrying a *Rosa26^LSL-YFP^* recombination reporter. Mice were infected with MNV and received tamoxifen by oral gavage at day 14 post-infection, thereby permanently labeling all cells expressing T-bet at the time of tamoxifen gavage with YFP **(Fig. 5a).** Mice were analyzed 3, 5, and 12 weeks after labeling to assess the persistence of tdTomato expression within YFP-labeled B cells **(Fig. 5a-b)**. YFP labelled B cell subsets were detected with the mLN, PP and the lamina propria of the small intestine and colon (**Extended data Fig. 5a-d)**. Within MNVP-specific MBC in PP and mLN, the vast majority of YFP^+^ MBC retained tdTomato expression for up to 12 weeks post-labeling (80-90% tdTomato+ of YFP labelled cells) **(Fig. 5c-d)**. Furthermore, we found stable T-bet expression regardless of the isotype B cells switch to, as tdTomato expression was observed in YFP^+^ IgG2c⁺ and IgA⁺ MNVP-specific MBC B cells in PP and mLN **(Fig. 5e-f)**. We next examined tdTomato expression in YFP-labeled B cells residing within the intestinal lamina propria. We found that the majority of YFP-labeled MBCs in both the small intestine and colon lamina propria, retained tdTomato expression over time **(Fig. 5g-h)**. In the small intestine, approximately 60% of YFP⁺ MBC retained tdTomato expression 12 weeks post-labeling, whereas retention remained above 80% in the colon **(Fig. 5g-h)**. Plasma cells showed limited maintenance of tdTomato expression. At 3 weeks post-labeling, approximately 40% of YFP⁺ plasma cells in the small intestine and ∼30% in the colon expressed tdTomato **(Fig. 5i-j)**. In the colon, this frequency was maintained up to 12 weeks post-labeling, whereas in the small intestine, tdTomato expression declined over time and stabilized at ∼20% of plasma cells by 12 weeks **(Fig. 5i-j)**. Because MNV-specific plasma cells constitute only a small fraction of the total plasma cell pool in the intestinal lamina propria, these values likely underestimate the persistence of T-bet expression within the antigen-specific plasma cell compartment. In addition, in the intestinal lamina propria antigen specific plasma cells or MBC could not be reliably identified using the MNVP probe, likely due to tissue processing interfering with probe binding (data not shown). Whereas T-bet expression in plasma cells was predominantly transient at both baseline and during infection, lamina propria MBC and secondary lymphoid organ B cell subsets exhibited sustained T-bet expression during MNV infection compared to homeostatic conditions **(Fig. 5c-h**, **Fig. 1i-j and Extended data Fig. 1f-g)**. Altogether, these data demonstrate that in contrast to homeostatic conditions, where T-bet expression is transient and associated with rapidly turning-over B cell populations (**Fig. 1i-j**), T-bet expression is durably maintained on MNVP-specific B cells for months following enteric viral infection.

**Figure 5.**
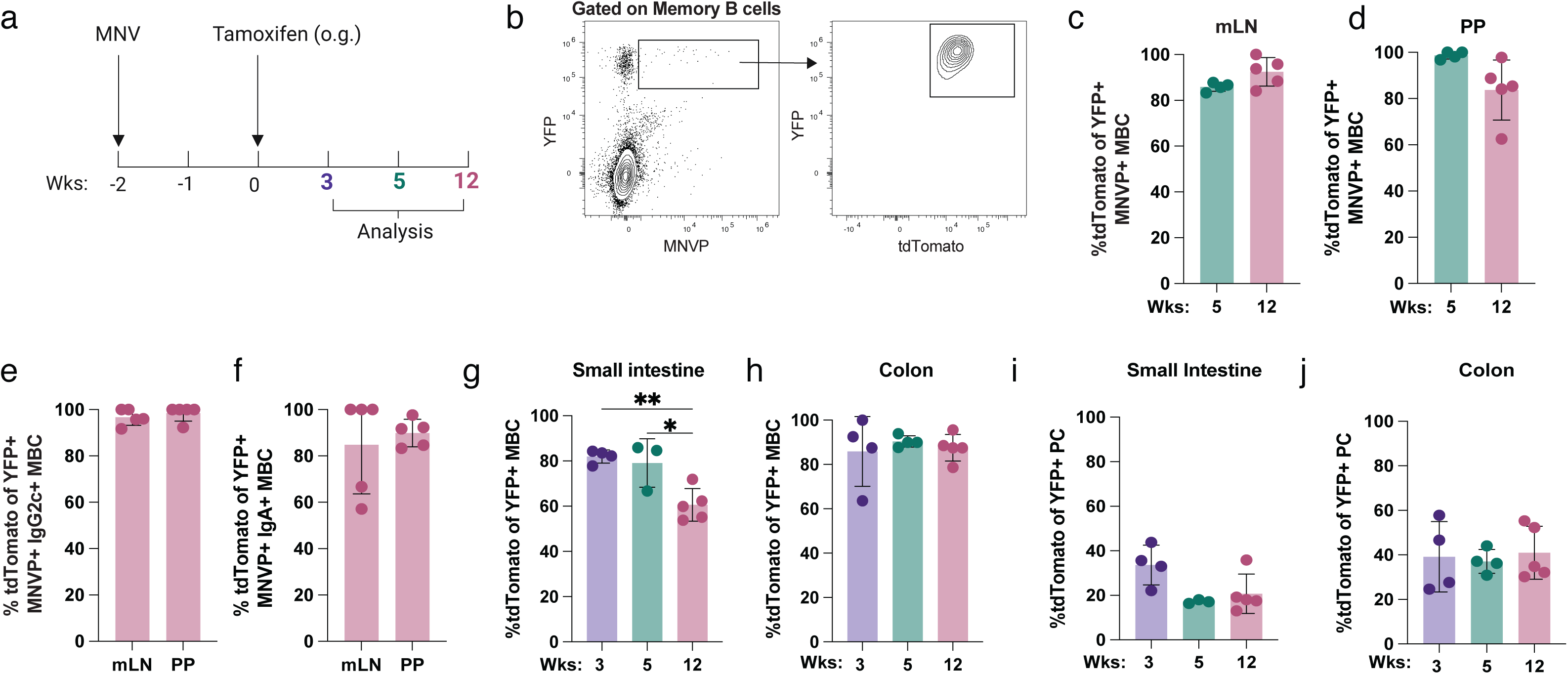
Enteric viral infection drives stable B cell T-bet expression. *Tbx21^tdTomato-creERT2^ Rosa26^LSL-YFP^* mice were infected with 1E6 PFU of MNV and administered tamoxifen 14 dpi by oral gavage, indicated organs were analyzed at indicated timepoints by flow cytometry. **(a)** Schematic for tracking stability of T-bet expression in B cells during MNV infection. **(b)** Representative flow cytometry plot showing gating strategy for tdTomato expression within YFP labelled MNVP+ cells **(c)** Frequency of tdTomato+ YFP+ MNVP+ memory B cells (MBC) in mLN. **(d)** Frequency of tdTomato+ YFP+ MNVP+ memory B cells (MBC) in PP. **(e)** Frequency of tdTomato+ YFP+ MNVP+ IgG2c-switched memory B cells (MBC) in mLN and PP. **(f)** Frequency of tdTomato+ YFP+ MNVP+ IgA-switched memory B cells (MBC) in mLN and PP. **(g)** Frequency of tdTomato+ YFP+ memory B cells (MBC) in small intestine lamina propria. **(h)** Frequency of tdTomato+ YFP+ memory B cells (MBC) in colon lamina propria. **(i)** Frequency of tdTomato+ YFP+ plasma cells (PC) in small intestine lamina propria. **(j)** Frequency of tdTomato+ YFP+ plasma cells (PC) in colon lamina propria. (**c-j**) Each dot represents a mouse, bars show mean, error bars show S.D; **(c-f)** Unpaired tests, **(g-h)** One-way ANOVA with Bonferroni correction for multiple comparisons. *P< 0.033(*), P< 0.002(**)*.

### T-bet is dispensable for IgA class-switching but required for effective GC responses during MNV infection

Canonically, T-bet is associated with class-switching to IgG2c^26^; however, our observation that T-bet is also expressed in IgA⁺ MNVP-specific B cells raised the question of whether T-bet is required for class-switching to IgA in this context. To address this, we examined the role of B cell-intrinsic T-bet in class-switching during MNV infection. To do so, we conditionally deleted *Tbx21* in B cells using *Tbx21^fl/fl^; CD23^Cre^* (*Tbx21^BΔ^*) mice and compared them to littermate control *Tbx21^fl/wt^; CD23^Cre^* (Ctrl) mice. We noted a significant decrease in the proportion of MNV-specific IgG2c⁺ GC B cells but no change in MNV-specific IgA⁺ GC B cells **(Fig. 6a, Extended data Fig. 6a)**. In line with these findings, *Tbx21^BΔ^* mice had a decrease in systemic MNV-specific IgG2c antibodies but no change in systemic or luminal MNV-specific IgA antibodies compared to infected littermate controls **(Fig. 6b-d)**. We also observed an increase in MNV-specific systemic IgG1, while no significant changes were detected in MNV-specific total IgG or IgG2b antibodies in *Tbx21^BΔ^* mice, a result consistent with prior studies showing that T-bet constrains class-switching to IgG1 **(Extended data Fig. 6b-d)**^26,51^. Together, these data show that within MNVP-specific B cells, T-bet is induced on both IgA⁺ and IgG2c⁺ cells and is selectively required for IgG2c class-switching but dispensable for IgA class-switching in response to MNV infection.

**Figure 6.**
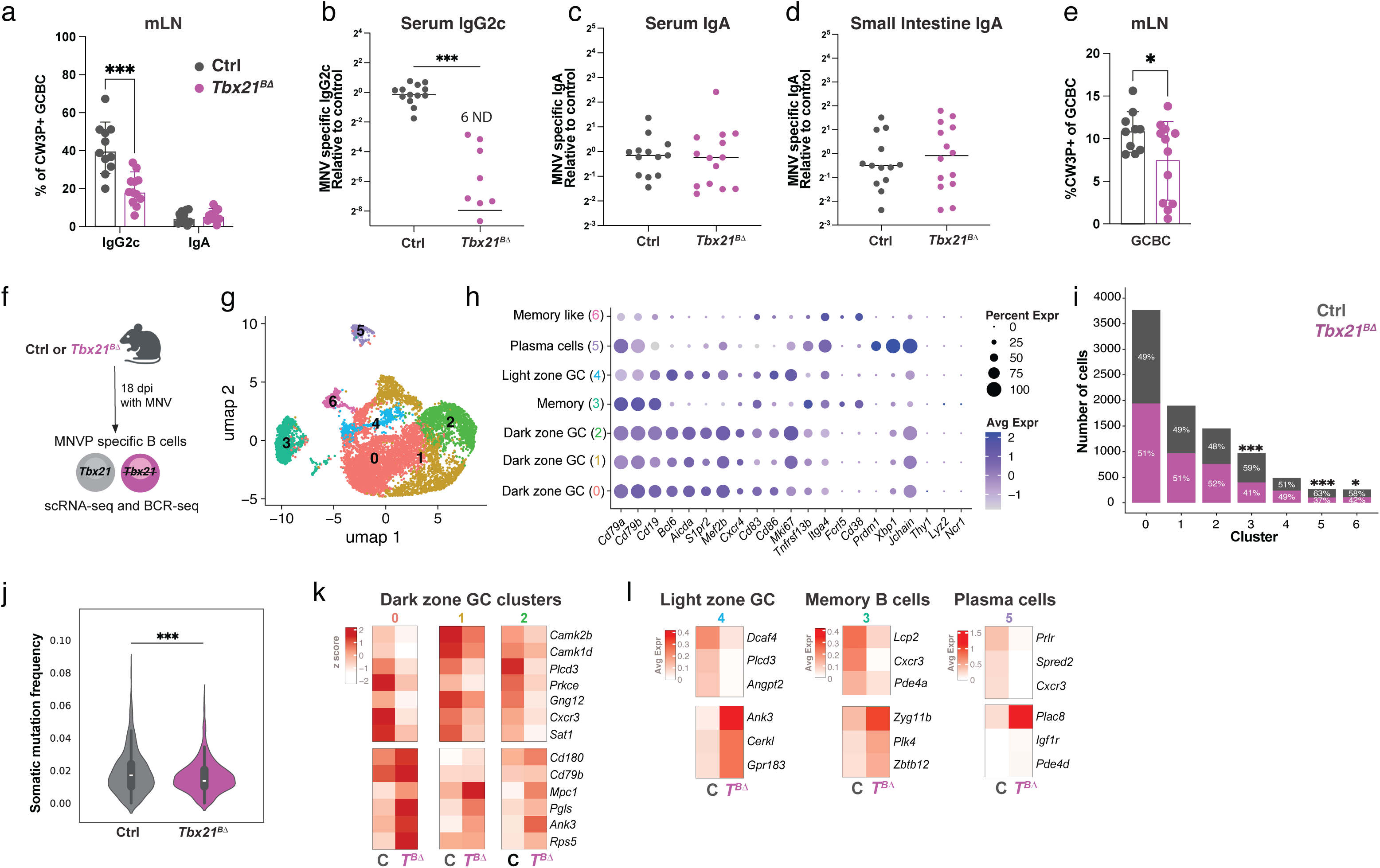
T-bet is dispensable for IgA class-switching but required for effective GC responses during MNV infection. **(a-c)** *Tbx21^fl/fl^ CD23^Cre^* (*Tbx21^BΔ^*) and control *Tbx21^fl/wt^ CD23^Cre^* (Ctrl) mice were infected with 1E6 PFU of MNV and analyzed 21 dpi by flow cytometry. **(a)** Frequency of MNVP+ IgG2c and IgA switched GC B cells (GCBC) in mLN. **(b-d)** MNV-specific antibodies were quantified by ELISA in serum and in small intestine luminal content at 21 dpi. **(b)** MNV specific IgG2c **(c)** and IgA in serum. Absorbance was normalized to control and displayed as fold change on log scale. **(d)** MNV specific IgA in luminal contents of small intestine. Absorbance normalized to control and displayed on log scale. **(e)** Frequency of MNVP+ GC B cells (GCBC) in mLN. **(f-l)** *Tbx21^fl/fl^ CD23^Cre^* (*Tbx21^BΔ^*) and control *Tbx21^fl/wt^ CD23^Cre^* (Ctrl) mice were infected with 1E6 PFU of MNV. Antigen specific (MNVP+) B cells were sorted 18 dpi and analyzed by scRNA-seq and BCR-seq. **(f)** Experimental Schematic. **(g)** UMAP visualization of scRNA-seq data from MNVP⁺ B cells colored by cluster identity following graph-based clustering. **(h)** Canonical marker genes used to define B cell subsets. Dot size indicates the proportion of cells expressing each gene and color intensity represents average log normalized expression. **(i)** Proportion and counts of cells derived from Ctrl and *Tbx21^BΔ^* within clusters. **(j)** Frequency of somatic hypermutation in non-naïve MNVP+ B cells (see methods for details). **(k)** Select differentially expressed genes between Ctrl and *Tbx21^BΔ^* cells across dark zone germinal center clusters (clusters 0-2). Values represent gene-wise z score scaled average expression across genotypes**. (l)** Select differentially expressed genes between Ctrl and *Tbx21^BΔ^* cells in memory (cluster 3), light zone GC (cluster 4), and plasma cell (cluster 5) populations. Values represent average log normalized expression within each cluster. **(a-e)** Each dot represents a mouse, bars show mean, error bars show S.D; **(a)** 2-way ANOVA with Bonferroni correction for multiple comparisons; **(b)** Mann-Whitney test; **(c-e)** Unpaired two-tailed t-test; **(i)** Hypergeometric test with Benjamini-Hochberg adjusted p-values; **(j)** Mann-Whitney test. *P < 0.033(*), P < .001(***)*.

Beyond class-switch recombination, expression of T-bet in all MNVP-specific B cells suggested that it may play a broader role in shaping B cell responses during viral enteric infection. Indeed, *Tbx21^BΔ^* mice showed reduced MNVP-specific GC B cells, suggesting altered GC B cell dynamics in the absence of T-bet **(Fig. 6e, Extended data Fig. 6e)**. To define the molecular programs downstream of T-bet that underlie these defects, we performed single-cell RNA sequencing (scRNA-seq) on MNVP-specific B cells isolated from mLN of *Tbx21^BΔ^* and littermate Ctrl mice, 18 days following MNV infection **(Fig. 6f)**. Unsupervised clustering of scRNA-seq data identified seven distinct B cell clusters **(Fig. 6g)**. Clusters were annotated based on canonical marker expression **(Fig. 6h)**. The largest clusters corresponded to GC B cells, with segregation into dark zone and light zone subsets with additional clusters including MBC, plasma cells, and a transitional memory-like population **(Fig. 6g-h)**. *Tbx21* deficiency did not broadly alter B cell subset identity, as we found no significant differences in compared cell identity module scores across genotypes, indicating that core subset identity is preserved in the absence of T-bet **(Extended data Fig. 6f)**. While most clusters contained comparable proportions of cells from *Tbx21^BΔ^* and Ctrl mice, MBC and plasma cells (clusters 3, 5, and 6) were significantly enriched for B cells originating from Ctrl animals, suggesting that T-bet is required for effective GC output and for the progression of MNV specific B cells into differentiated states **(Fig. 6i).** In line with this, BCR-seq revealed reduced somatic hypermutation within MNV-specific B cells from *Tbx21^BΔ^* mice within the non-naive B cell compartment, defined as cells with at least 2% divergence from germline, compared with controls, further supporting a role for T-bet in proper B cell maturation within the GC **(Fig. 6j).**

We next performed differential expression analysis comparing *Tbx21^BΔ^* and Ctrl MNV-specific B cells across clusters and found that T-bet regulates distinct gene programs that support activation, selection, and differentiation in the GC B cell response and progression of B cells to differentiated states (**Fig. 6k-l, Table 1**). In particular, deletion of *Tbx21* in B cells altered gene expression programs associated with cellular metabolism *(Prlr* and *Sat1)*, as well as with BCR and cAMP signaling (like *Plcd3, Lcp2, Pde4a, Cd79b,* and *Camk2b*) (**Fig. 6k-l)**. Cellular positioning is also essential for mounting an effective B cell response, encompassing both the spatial organization of cells within secondary lymphoid organs and their migration to effector sites^22^. In the GC, B cells transition between the dark zone, where they undergo clonal expansion and somatic hypermutation, and the light zone, where antigen acquisition and T follicular helper cell mediated selection occur^52^. Disruption of this coordinated positioning can impair germinal center output^16,52^. Consistent with defects in *Tbx21^BΔ^* cells, *Gpr183*, which encodes the chemokine receptor EBI2, was increased in light zone GC B cells (**Fig. 6l)**. As EBI2 downregulation is required for B cells to localize appropriately within the follicle and efficiently participate in germinal center responses, these data suggest that T-bet deficient MNV specific GC B cells may fail to properly regulate follicular positioning, thereby contributing to impaired GC function^53,54^. Beyond organization and signaling within the GC, cellular positioning also governs the migration of differentiated B cells to peripheral effector sites. In this context, *Cxcr3* was dysregulated in several B cell populations when *Tbx21* was deleted, suggesting that T-bet sustains a conserved chemokine receptor signaling axis that may support cellular positioning during enteric viral infection. Given the reduced representation of *Tbx21^BΔ^* cells within memory and plasma cell clusters, these data suggest that T-bet may support not only GC organization but also the trafficking or accumulation of post GC effector populations during enteric viral infection **(Fig. 6k-l).** Indeed, prior studies suggest that *Cxcr3* supports the accumulation of IgA secreting B cell populations in mucosal tissues during influenza and rotavirus infection^18,47,55^. The combination of these transcriptional changes likely underlie the reduced GC B cell numbers and diminished somatic hypermutation observed in the absence of T-bet (**Fig. 6e, j)**, as well as the decreased functional output marked by reduced representation of MBC and plasma cells (**Fig. 6i).** Together, these findings suggest that T-bet sustains the signaling and metabolic fitness required for MNV specific B cells to efficiently participate in GC responses and progress into downstream effector states.

### T-bet⁺ B cells are required for circulating and mucosal antibodies providing protection to reinfection

Given the expansion and stable maintenance of T-bet expression in B cells during infection, we next sought to determine how loss of this B cell subset would impact antiviral immunity and whether these cells are required for protective responses against re-encounter with the pathogen. To test the role of T-bet-expressing B cells in enteric antiviral responses, we used *Pax5^fl/fl^; Tbx21^tdTomato-cre^* mice, which have selective ablation of T-bet⁺ B cells via deletion of *Pax5*, a transcription factor essential for B cell identity and maintenance^56^. In *Pax5^fl/fl^; Tbx21^tdTomato-cre^*mice, we observed a marked reduction in T-bet expressing B cells within lymphoid organs as well as within the intestinal lamina propria compared to littermate controls **(Extended data Fig. 7a-f)**. Notably, ablation of T-bet⁺ B cells did not alter the proportion of total B cells or most major immune populations, except for a minor increase in TCRβ⁺ T cells and eosinophils in the small intestinal lamina propria, which may reflect that loss of T-bet⁺ B cells may subtly reshape the local immune landscape without broadly disrupting overall immune composition **(Extended data Fig. 7g-j)**. We next evaluated the contribution of T-bet-expressing B cells in response to norovirus infection. Ablation of T-bet⁺ B cells resulted in a marked reduction in MNVP-specific GC B cells **(Fig. 7a-b)** and MBC **(Extended data Fig. 7k-l)**, indicating that T-bet⁺ B cells comprise the majority of the virus-specific B cell response during MNV infection, with no other B cells able to compensate for their loss. Consistent with a loss of MNV-specific B cell response, ablation of T-bet expressing B cells led to near complete loss of MNV-specific IgG antibodies in circulation, most prominently IgG2c, in *Pax5^fl/fl^; Tbx21^tdTomato-cre^* mice compared to littermate controls **(Fig. 7c-d).** In addition, *Pax5^fl/fl^; Tbx21^tdTomato-cre^* mice had a near complete loss of MNV-specific IgA antibodies in the serum as well as in the luminal contents of the small intestine and colon, with antibody levels at or below the limit of detection. **(Fig. 7e-g).** Importantly, we found no change in the establishment of MNV infection in *Pax5^fl/fl^; Tbx21^tdTomato-cre^* mice 3 days following the primary infection, indicating that the differences observed were not the result of MNV failing to infect T-bet⁺ B cell-deficient mice **(Extended data Fig. 7m)**. Overall, these data indicate that T-bet expressing B cells constitute most of the MNV-specific B cell response and ablation of T-bet expressing cells leads to a near complete loss of MNV-specific systemic and mucosal antibody responses.

**Figure 7.**
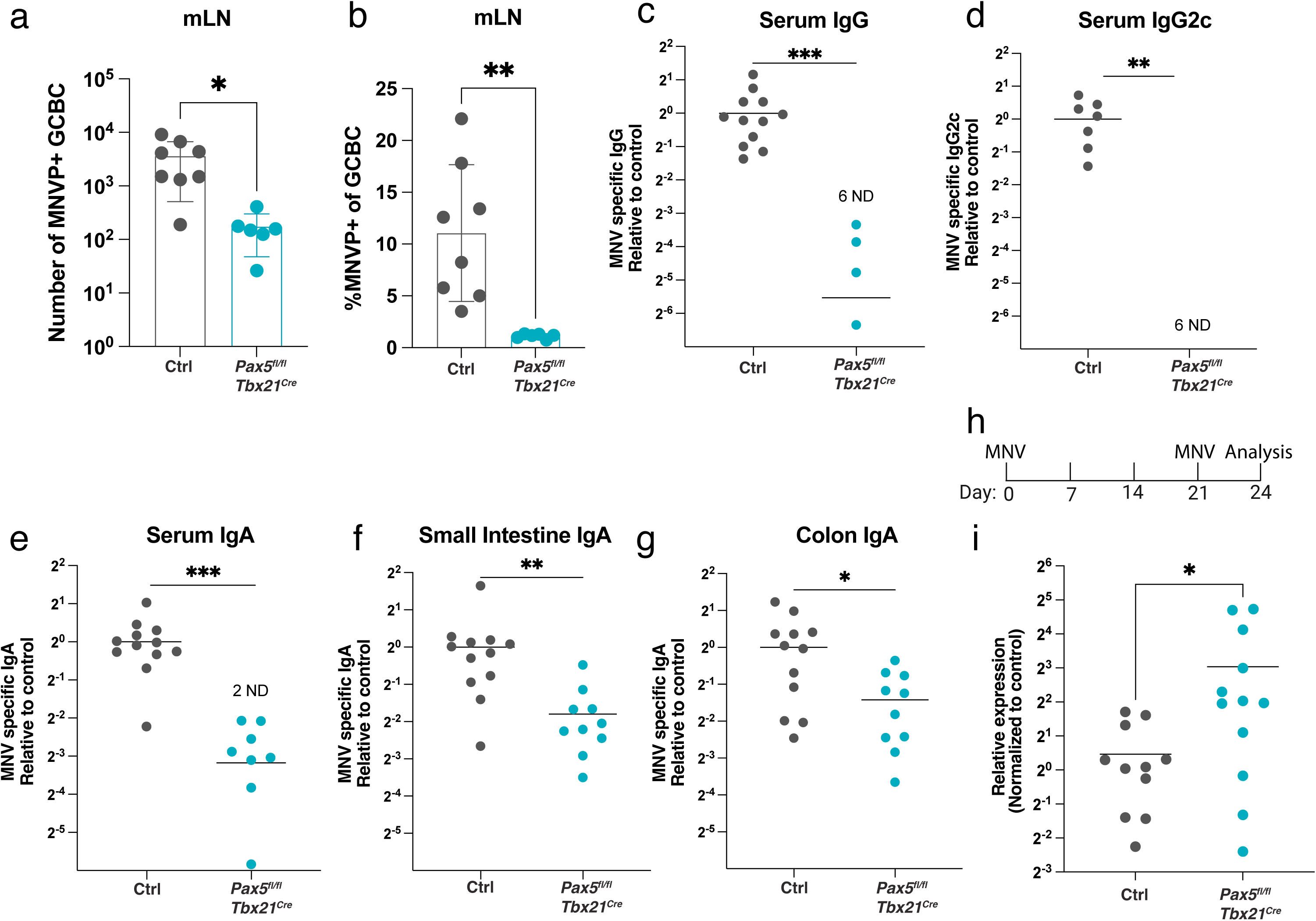
T-bet⁺ B cells are required for circulating and mucosal antibodies providing protection to reinfection. **(a-g)** *Pax5^fl/fl^ Tbx21^Cre^* and control *Pax5^fl/wt^ Tbx21^Cre^* (Ctrl) mice were infected with 1E6 PFU of MNV and analyzed at 21dpi by **(a-b)** flow cytometry and **(c-g)** ELISA. **(a)** Number and **(b)** frequency of MNVP+ GC B cells (GCBC) in mLN. **(c-g)** MNV-specific antibodies **(c)** IgG in serum **(d)** IgG2c in serum **(e)** IgA in serum **(f)** IgA in small intestine luminal contents and in **(g)** colon luminal contents. Absorbance was normalized to control and displayed on log scale. **(h-i)** *Pax5^fl/fl^ Tbx21^Cre^* and control *Pax5^fl/wt^ Tbx21^Cre^* (Ctrl) mice were infected with 1E6 PFU of MNV. 3 weeks post infection mice were re-infected with MNV. Quantification of viral RNA from distal colon tissue was performed using RT-qPCR and normalized to intestinal actin expression 3 days post re-infection. **(h)** Schematic of viral infection and analysis. **(i)** Relative MNV RNA abundance in the distal ileum at day 3 post challenge, determined by RT-qPCR. Data are presented as fold change relative to controls, calculated using the 2^-ΔΔCT^ method with Actb as the internal reference gene. **(a-i)** Each dot represents a mouse, bars show mean, error bars show S.D.; Unpaired two-tailed t-test, *0.033(*), P 0.002(**), P < .001(***)*.

Memory B cell and antibody responses are hallmarks of adaptive immunity. As ablation of T-bet⁺ B cells selectively impaired these responses, we examined whether T-bet expressing B cells are required for protection against MNV re-infection. To test this, we used a re-challenge model in which *Pax5^fl/fl^; Tbx21^tdTomato-cre^* mice and littermate controls were infected with MNV and subsequently rechallenged with virus three weeks later, following resolution of the primary infection **(Fig. 7h)**. We found that *Pax5^fl/fl^; Tbx21^tdTomato-cre^* mice displayed significantly higher viral RNA levels in the distal ileum 3 days after viral re-challenge compared to littermate controls indicating impaired protective immunity in the absence of T-bet expressing B cells **(Fig. 7i)**. These findings demonstrate that T-bet-expressing B cells critically contribute to protection against future MNV infection. Altogether, these data indicate that T-bet-expressing B cells are a distinct subset of B cells responsible for both systemic IgG and mucosal IgA antibody production against an enteric virus and are necessary for protection against subsequent infection.

Together, these findings identify T-bet as a central regulator in the humoral immune response to enteric viral infection. MNV infection induces a sustained T-bet transcriptional program across antigen-specific B cell subsets that integrates germinal center dynamics, cellular signaling, and tissue positioning to support antiviral immunity. Although IgG2c⁺ and IgA⁺ B cells arise from distinct progenitors, both populations express T-bet. Thus, T-bet functions not simply as a regulator of class switching, but as a broader organizer of antiviral B cell responses during enteric viral infection.

## DISCUSSION

Understanding how B cells coordinate mucosal defense is particularly important given the rising incidence of intestinal inflammatory disorders^57^. The intestine represents the dominant site of antibody production in mammals, reflecting a substantial energetic investment in barrier protection^3^. This scale of humoral activity implies the existence of specialized B cell populations adapted to distinct functional roles. Indeed, tissue-specific specialization has been described for multiple immune cell types, including macrophages, regulatory T cells, and plasma cells, suggesting that local environmental cues shape distinct functional states within the immune system^5,58,59^. In this context, we find that T-bet-expressing B cells are a distinct subset of B cells responsive to microbial cues, and dynamically maintained. Interestingly, we found a marked difference in the stability of T-bet expression in the absence or presence of infection, indicating that inflammatory cues in response to infection may reinforce and stabilize T-bet expression to induce a durable population.

In response to infection, T-bet-expressing B cells constitute a dominant component of the antiviral response. During MNV infection, we observe robust and stable induction of T-bet in the majority of virus-specific B cells, and these cells are required for the production of virus-specific systemic IgG and luminal IgA antibodies. Interestingly, we find that B cells expressing the same transcription factor can undergo class-switching to either IgA or IgG2c. Analysis of BCR repertoires from MNV-reactive B cells further suggests that these outcomes arise largely in parallel, as we observe minimal clonal overlap between IgG2c⁺ and IgA⁺ populations. This raises the possibility that antigen-experienced B cells are differentially exposed to distinct, tightly regulated environmental cues, such as IFNγ or TGFβ, that bias the direction of class switching^26,60,61^. Notably, this process does not appear to be determined solely by the anatomical site of induction as both PP and mLN contain IgA⁺ and IgG2c⁺ B cells, albeit at different frequencies. This observation is consistent with previous work suggesting that cytokine signals governing class-switching are spatially restricted and locally regulated within lymphoid tissues^22^. Consistent with this idea, the limited clonal overlap between IgG2c⁺ and IgA⁺ repertoires argues against sequential switching between these isotypes during MNV infection and instead supports the notion that distinct signals independently induce class switch recombination in T-bet expressing B cells.

Furthermore, stable induction of T-bet during MNV infection suggested functions extending beyond class switching. Consistent with this, scRNA-seq revealed that T-bet deletion in B cells reshaped transcriptional programs linked to cellular migration and positioning, intracellular signaling, and metabolic support in MNVP-specific B cells. These effects were most evident within GC subsets, where T-bet deficiency also reduced somatic hypermutation, indicating compromised selection efficiency and germinal center quality. This is consistent with malaria infection, where T-bet is important for the quality of germinal center responses by promoting commitment to the dark zone and supporting affinity maturation^16^. In addition, our data suggest that T-bet may also regulate post GC effector differentiation and trafficking as *Cxcr3* was reduced in T-bet deficient MNV-specific B cells and memory and plasma cell clusters were enriched for control cells, suggesting delayed progression into these states. This is consistent with prior work showing that *Cxcr3* supports the accumulation of IgA secreting B cell populations in mucosal tissues and that T-bet expression controls antibody secreting cell recall responses later in the course of infection^18,19,47,55^. Thus, T-bet expression within B cells tunes the functional fitness of adaptive responses and may explain why stable T-bet expression is maintained after enteric viral infection.

Noroviruses remain the leading cause of viral gastroenteritis worldwide, and no licensed vaccines are currently available^62^. Our findings identify T-bet-expressing B cells as essential for antiviral mucosal immunity, with implications for mucosal vaccine development. T-bet expression in B cells has also been associated with autoimmunity. During intestinal inflammation, Intestinal Bowel Disease-associated SNPs are enriched at T-bet binding sites and circulating T-bet⁺ B cell frequencies correlate with Crohn’s disease severity, suggesting context-dependent roles for this subset^63,64^. Because viral infection induces sustained T-bet expression, chronic activation of this program in inflammatory settings may contribute to dysregulated responses^65^. Thus, defining the signals that drive T-bet⁺ B cell differentiation in the intestine may therefore inform strategies to promote protective immunity while limiting maladaptive inflammation.

## MATERIALS AND METHODS

### Mice

*Tbx21^tdTomato-^*^T2Acre^ and *Tbx21^tdTomato-T2A-creERT2^*mice were described previously^32^. *Pax5^f^* (JAX stock #028100), *Tbx21^f^ (*JAX stock #022741, originally developed by the laboratory of Steve Reiner), *CD23^cre^* (JAX stock #028197) and *Rosa26^Lox-STOP-Lox-YFP^* (JAX stock #006148) mice were purchased from the Jackson Laboratory^56,66,67^. Mice were 6 to 10 weeks old at the time of analysis, except for aging experiments where mice were 10 to 26 weeks old. Male and female mice were used depending on availability, as sex did not appear to have a major impact on the measurements made. In all cases, mice were compared with littermate controls. Generation and treatment of mice were performed under protocol 23-0004-1 approved by Scripps Research Institutional Animal Care and Use Committee. All mouse strains were maintained in the animal facility at TSRI in specific pathogen–free (SPF) conditions in accordance with institutional guidelines and ethical regulations.

### MNV infections

Murine leukemic monocyte–macrophage RAW264.7 cells and human embryonic kidney 293T cells (293T) were purchased from ATCC and maintained in Dulbecco’s modified Eagle’s medium (DMEM) supplemented with 10% (v/v) fetal bovine serum and 1% penicillin–streptomycin. Murine norovirus (MNV-CW3) was provided by Ken Cadwell (University of Pennsylvania) and propagated in RAW264.7 cells. Concentrated viral stocks were prepared as previously described^68^. Briefly, supernatants from 293T cells transfected with plasmids encoding the viral genome were used to infect RAW264.7 cells to amplify virus production. Viral particles were concentrated by ultracentrifugation using an SW28 rotor and resuspended in endotoxin-free PBS. Stock viral titers were determined by plaque assay, as described previously^69^. For *in vivo* infections, mice were orally gavaged with 10⁶ plaque-forming units (PFU) of virus in 150 µL of PBS, or mock treated with PBS alone. MNV viral RNA was quantified by RT-qPCR using RNA extracted from fecal samples and a 1-inch segment of distal ileum. Viral RNA was isolated using the Qiagen RNA isolation kit according to the manufacturer’s instructions and reverse transcribed into cDNA using NEB reverse transcription reagents (LunaScript). qPCR was performed using MNV specific primers described previously^70^. For distal ileum samples, viral RNA was measured by relative quantification using the 2^-ΔΔCT method with murine *Actb* as the internal reference.

### Tamoxifen administration

Tamoxifen (40 mg/ml) was dissolved in corn oil (Sigma-Aldrich). Mice were orally gavaged with one dose of 200uL of tamoxifen emulsion.

### Antibiotic treatment

Antibiotic treatment was performed as previously described^71^. Briefly, streptomycin (5 mg/mL), colistin sulfate (1 mg/mL), and ampicillin (1 mg/mL) were dissolved in mouse drinking water and filter sterilized. Experimental mice received antibiotic treated water, while control mice received sterile drinking water without antibiotics, for 14–18 days prior to analysis.

### MNVP probe

The P domain of VP1 from MNV strain CW3 (residues 225–541) was cloned into a pET-21a expression vector. The resulting construct was co-transformed with a BirA expression plasmid (pBirAcm) into *Escherichia coli* BL21 cells. Cultures were grown at 37°C to mid-log phase, cooled to room temperature on ice, and protein expression was induced overnight at 16°C with 0.2 mM IPTG in the presence of 20 µM biotin. Cells were harvested and lysed by sonication in lysis buffer containing 50 mM Tris-HCl (pH 7.5), 150 mM NaCl, 0.1% SDS, protease inhibitors (Roche), and benzonase. Lysates were clarified by centrifugation, and MNVP protein was purified from the supernatant using Ni–nitrilotriacetic acid (Ni-NTA) affinity chromatography (Qiagen), followed by 3 washes with an imidazole gradient (20mM, 40mM, 60mM) and elution with 300 mM imidazole. Eluted protein was concentrated, dialyzed overnight at 4°C into PBS, and further purified by size exclusion chromatography. Protein expression and biotinylation were confirmed by western blotting using anti-His antibodies and streptavidin-HRP. Fluorescent MNVP probes were generated as previously described^72^. Briefly, biotinylated MNVP was tetramerized with fluorophore-conjugated streptavidin (APC or BV421) at a 4:1 molar ratio of protein to streptavidin. Streptavidin was added stepwise in ten equal additions, with 10-minute incubations at room temperature in the dark between each addition. Tetramerized probes were stored at −80°C in 50% glycerol and aliquots were maintained at 4°C during experimental use.

### Microscopy

Confocal imaging was performed under standard conditions as previously described^22^. Briefly, mLN were fixed in 4% paraformaldehyde for 2 h at room temperature, followed by dehydration in 30% sucrose in PBS at 4°C. Tissues were then embedded in OCT compound (Sakura Tissue-Tek), snap-frozen, and sectioned at 10 μm. Sections were fixed in acetone for 20 min at −20°C, air dried overnight, and rehydrated in PBS for 10 min prior to staining. Staining was performed overnight in 1X PBS containing 10% donkey serum (Jackson ImmunoResearch) and 0.3% Triton X-100. Sections were mounted in SlowFade mounting medium (Life Technologies) and imaged on an LSM880 confocal microscope (Carl Zeiss) using a 40X oil immersion objective. Images were processed and analyzed in ImageJ [version 2.0.0-rc-68/1.52g; National Institutes of Health (NIH)].

### Cell isolation for flow cytometry

For mLN and PP, lymphocytes were isolated by mechanical disruption using a 1mL syringe plunger and filtration through a 100-μm cell strainer in staining buffer (2% heat inactivated fetal bovine serum (FBS), 1 mM EDTA, 10 mM Hepes, 1X PBS). For colonic lymphocyte isolation, the cecum was removed, and the colon was dissected free of fat, opened longitudinally, and washed extensively in 1X PBS to remove luminal contents. Tissue was cut into 1–2 cm pieces and incubated in wash medium containing 5 mM EDTA and 1 mM dithiothreitol in a 50 mL screw cap tube, shaking horizontally at 250 RPM for 15 minutes at 37 °C. Following brief vortexing, epithelial and intraepithelial cells were removed by filtration through a 200μm cell strainer. Remaining tissue was washed once more in PBS and filtered again. Tissue was then digested in warm wash medium supplemented with 0.2 U/mL collagenase type 1 and 1 U/mL DNase I in the presence of ceramic beads, shaking at 250 RPM for 35 minutes at 37 °C. The resulting suspension was filtered through a 100μm strainer, centrifuged to remove debris and enzyme, washed, and resuspended in 40% Percoll prior to centrifugation. Small intestine samples were processed using the same protocol as the colon, with the following modifications: PP were removed, epithelial cells were depleted by an additional shaking step using 5 mM EDTA in PBS at 37 °C, the final digestion volume was 30 mL, and collagenase concentration was increased 1.5-fold. Wash medium contained the following components: 1X RPMI 1640, 2% FBS, 10mM HEPES Buffer (Corning CellGro), 1% Penicillin-Streptomycin (Corning CellGro), 1% L-Glutamine (Corning CellGro).

### Flow cytometry

All centrifugation steps were performed at 900 g for 3 min at 4°C. All staining was carried out in 96-well V-bottom plates in a final volume of 50 μL. Cells were first stained with viability dye and Fc receptors were blocked in PBS for 10 min at 4°C. Cells were then washed with 150μL flow staining buffer (PBS containing 0.2% FBS, 2 mM EDTA, and 1% penicillin-streptomycin), followed by extracellular staining in flow staining buffer for 30 min at 4°C. After staining, cells were washed with 150 μL flow staining buffer and resuspended in 200 μl flow staining buffer. Samples were passed through a 100μm mesh before acquisition.

For intracellular Blimp1 staining, cells were stained at 4°C in 1× Perm/Wash buffer. Samples were then washed twice with 200 μL 1X Perm/Wash buffer, resuspended in 200 μL staining buffer, and passed through a 100 μm nylon mesh. All samples were acquired on an Aurora cytometer (Cytek Biosciences) and analyzed using FlowJo software (BD Biosciences).

### Antibodies

**The following antibodies were used for flow cytometry experiments:** B220 (RA3-6B2), CD90.2 (53-2.1), CD19 (6D5), CD45 (30-F11), IgD (11-26c.2a), CD95/Fas (Jo2 and SA367H8), TCRb (H57-597), CD11b (M1/70), Live Dead Blue viability dye (Thermofisher, L23105), CD38 (90), GL7 (GL7), IgG2b (R12-3), IgG1 (X56), IgG2c (Southern Biotech, 1079-05), IgA (mA-6E1), CD64 (X54-5/7.1), SiglecF (E50-2440), NK1.1 (PK136), Gr-1 (RB6-8C5), FeERR1 (46082), CD11c (N418), TCRgd (GL3), LY6C (HK1.4), cKit (2B8), and CD44 (IM7).

**The following antibodies were used for microscopy experiments:** DsRed (rabbit polyclonal, Takara; and goat polycloal, MyBiosource), B220 (RA3-6B2), IgD (11-26c.2a).

### Enzyme-linked immunosorbent assay (ELISA)

For detecting MNV specific antibodies, whole lysed virus was used. Briefly, virus was resuspended in PBS containing 0.2% Triton X-100 and subjected to two freeze–thaw cycles. Lysed virus was quantified by Bradford assay and coated onto ELISA plates at 2 µg/mL in carbonate coating buffer overnight at 4°C. Plates were blocked with 1% bovine serum albumin in 1× PBS for 2 hours at 37°C. Serum samples were diluted in ELISA buffer and incubated on plates for 1 hour at 4°C. Intestinal luminal samples were resuspended in 1X PBS containing protease inhibitors (small intestine, 1 mg/mL; colon, 0.3 mg/mL), vortexed, and clarified by sequential centrifugation at 400 g for 5 minutes followed by 8000 g for 5 minutes. Only the clear supernatant was collected and either used immediately or flash frozen and stored at −80°C. Prior to use, luminal samples were diluted in ELISA buffer to final concentrations of 0.5 mg/mL for small intestine samples and 0.08 mg/mL for colonic samples. HRP conjugated secondary antibodies against total IgG, IgG2c, IgG1, and IgG2b (SouthernBiotech) were used for detection. IgA was detected using biotinylated anti IgA followed by streptavidin HRP. Background signal obtained with secondary antibody alone was subtracted from all measurements. For ELISAs using MNVP a similar protocol was followed with the exception that plates were first coated with 1 ug/uL of anti-his antibody in carbonate coating buffer overnight at 4°C. 2ug/mL of MNVP was subsequently added for 1 hour at 4C.

### Single cell RNA sequencing (scRNA-seq)

An equivalent number of Ctrl and *Tbx21^BΔ^* MNVP-specific GC B cells and MBC were sorted using the MNVP probe according to the gating strategy shown in **(Fig. 1a)**. A total of 12,000 cells were used for library preparation with the 10x Genomics Single Cell 5′ Reagent Kit v2 according to the manufacturer’s instructions. BioLegend TotalSeq C hashtag oligonucleotides were used to label Ctrl and *Tbx21^BΔ^* samples, which were then pooled and loaded into the same sequencing lane. Libraries were sequenced on the Element AVITI platform. Sequencing reads were processed, aligned to the mm10 2020 reference genome, and demultiplexed using Cell Ranger v9.0.1 with default parameters. Hashtag oligonucleotide reads were also processed and demultiplexed using Cell Ranger with default parameters. Downstream analysis was performed in R using a custom script built with the Seurat package v5.4.0.

After alignment and demultiplexing, RNA and hashtag oligonucleotide matrices were matched using shared cell barcodes. Cells were assigned to Ctrl or *Tbx21^BΔ^* samples using the Seurat HTODemux function following centered log ratio normalization of hashtag oligonucleotide counts, and only singlet cells were retained for downstream analysis. Quality control filtering removed cells with fewer than 200 or more than 5,000 detected genes or greater than 20% mitochondrial transcripts. RNA expression values were log normalized and 2,000 highly variable genes were identified using the vst method. Principal component analysis was performed on variable genes, and the first 10 principal components were used to construct a shared nearest neighbor graph for clustering. Cells were clustered using Seurat, and cluster identities generated at resolution 0.1 were used for downstream analyses.

Dot plots displaying gene expression across clusters were generated using the Seurat DotPlot function, which visualizes the average log normalized expression of each gene together with the proportion of cells expressing that gene within each cluster. Bar plots showing the distribution of Ctrl and *Tbx21^BΔ^* cells across clusters were generated in R using ggplot2 based on cell counts per cluster. To evaluate Ctrl cell enrichment within individual clusters relative to the global dataset baseline, a hypergeometric test was performed using the phyper function in R. The background population comprised all Ctrl and *Tbx21^BΔ^* cells. Resulting p-values were adjusted for multiple comparisons using the Benjamini-Hochberg procedure to control the false discovery rate.

Heatmaps were generated to visualize transcriptional differences between Ctrl and *Tbx21^BΔ^* cells across clusters. Differential gene expression between genotypes was first performed independently within each cluster using Seurat FindMarkers with the Wilcoxon rank sum test and an initial log2 fold change threshold of 0.1. Genes encoding immunoglobulin variable regions (Igkv, Ighv, Iglv), predicted genes beginning with “Gm”, and genes ending in “Rik” were excluded from downstream analyses.

For dark zone germinal center clusters (clusters 0–2), differentially expressed genes were filtered using an adjusted p value < 0.05 and an absolute log2 fold change ≥ 0.1. Average expression values for Ctrl and *Tbx21^BΔ^* cells within each cluster were calculated using Seurat AverageExpression, and gene expression values were z score scaled across genes prior to visualization. Heatmaps therefore display relative gene expression differences between genotypes across clusters.

For visualization of transcriptional programs within clusters 3-6, differential expression tables were further filtered to retain genes expressed in at least 5% of cells in either genotype and with an absolute log2 fold change > 0.25. Genes were ranked by fold change and the top 25 genes per cluster were selected for visualization. These heatmaps display average log normalized expression values for Ctrl and *Tbx21^BΔ^* cells within each cluster without additional scaling. Heatmaps were generated using the ComplexHeatmap R package, and only representative genes meeting the above criteria are shown in the final figures.

Cluster-specific gene modules were defined using the top 100 upregulated markers (ranked by p-value) derived exclusively from Ctrl cells via Seurat’s FindAllMarkers function (Wilcoxon Rank Sum test, logfc.threshold = 0.25, min.pct = 0.25). To evaluate transcriptional preservation, these WT-derived modules were scored across the entire dataset (Ctrl and *Tbx21^BΔ^* cells) using Seurat’s AddModuleScore function. Statistical comparisons of the resulting matched module scores between Ctrl and *Tbx21^BΔ^* cells within individual clusters were performed using a Wilcoxon Rank Sum test.

### B cell receptor sequencing (BCR-seq)

For clonotype analysis, in an equivalent number of MNVP-specific IgA-switched and IgG2c-switched GC B cells and MBC were sorted and pooled from mLN of 6 *Tbx21^tdTomato-^*^T2Acre^ mice 14 days post infection using the gating strategy shown in **(Fig. 1a)**. For detecting somatic hypermutation frequencies, an equivalent number of MNVP-specific GC B cells and MBC were pooled and sorted from mLN of 5 Ctrl and 4 *Tbx21^BΔ^* mice 18 days post infection. VDJ library preparation was performed with the 10x Genomics Single Cell 5′ Reagent Kit v2 according to the manufacturer’s instructions. Libraries were sequenced on the Element AVITI platform. Sequencing reads were processed, aligned to the mm10 2020 reference genome, and demultiplexed using Cell Ranger v9.0.1 with default parameters. B cell receptor (BCR) sequences were subsequently annotated using the abstar toolkit, as described previously^73^. From these data, productive heavy-light chain BCR pairs were identified. During preprocessing, only observations containing exactly one heavy chain and one light chain sequence were retained. Clonal lineages were inferred using the clonify function in the scab toolkit^74^. Somatic hypermutation was evaluated using the abstar toolkit, by computing sequence identity relative to murine germline V gene segments^73^. Mutation frequency was defined as 1 – v_identity. To prevent the low background mutation rate of naïve cells from obscuring treatment-associated differences in mutation frequency, cells were stratified into naïve and non-naïve populations. Naïve cells were defined as IgM-expressing cells with ≥98% similarity to germline V gene sequences (v_identity ≥0.98).

Statistical comparisons of mutation frequency between groups were performed using the Mann-Whitney U test. Source code is available upon request.

### Statistical Analysis

All statistical tests are indicated in the corresponding figure legends. All statistical analyses, except those related to scRNA-seq, were performed using GraphPad Prism 6. Statistical analyses for scRNA-seq were performed in R. Data were assumed to be normally distributed, although this was not formally tested. All experiments were repeated at least twice. P values were considered significant as follows: *P< 0.033(*), P< 0.002(**). P < .001(***).* Numbers of mice and measures of central tendency and variation are indicated in the corresponding figures.

## Extended Data

**Table 1:** List of DE genes between Ctrl and *Tbx21^BΔ^* cells within each cluster of B cells identified using scRNA-seq.

**Extended Data Figures 1-7.**

## Acknowledgments

We thank K. Cadwell for providing the murine norovirus strain, and all the members of the Mendoza laboratory for technical input and discussion.

## Author contributions

A.M. and A.M. designed the experiments, interpreted the data, and wrote the manuscript. A.M. performed experiments and analyzed data with help from T.B., A.W., S.D., K.F., D.M., S.M., N.W. S.D and D.M contributed to data analysis. B.B, M.Z. and H.H. interpreted data.

## Data and materials availability

The accession number for the scRNA-seq datasets reported in this paper will be accessible through GEO and made available upon publication. All other data needed to evaluate the conclusions in the paper are present in the paper or the Extended Data.

---

**Extended data Figure 1.**
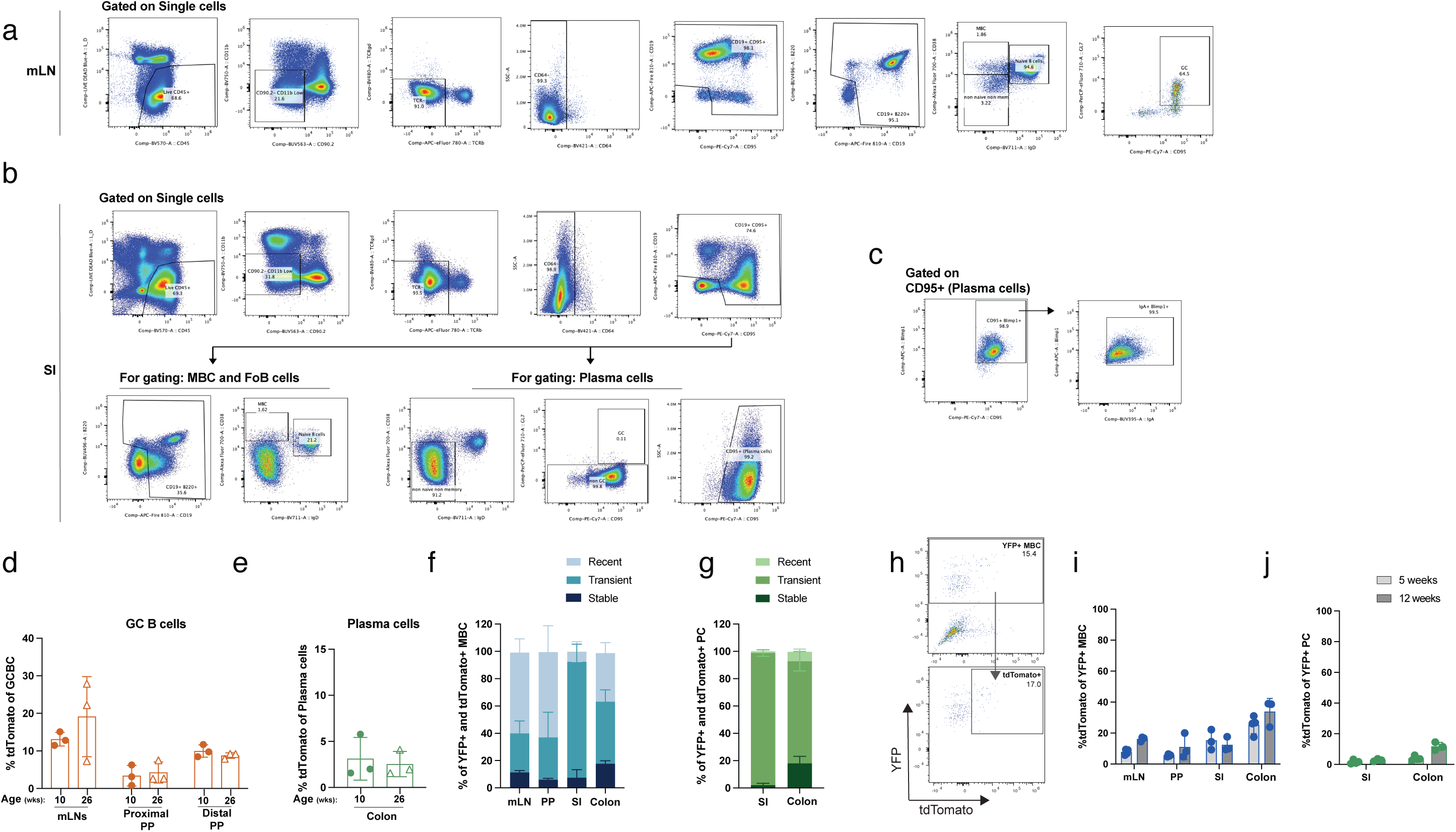
**(a-e)** Uninfected *Tbx21^tdTomato-cre^* mice were analyzed by flow cytometry. **(a)** Gating Strategy to identify B cell subsets within PP and mLN. (**b**) Gating Strategy to identify B cell subsets within lamina propria. **(c)** Validation of CD95 as a plasma cell marker using intracellular Blimp1 staining in lamina propria. **(a-c)** Mice were analyzed at 8 weeks of age. **(d)** Frequency of tdTomato+ GC B cells within mLN of 10- and 26-week-old mice. **(e)** Frequency of tdTomato+ Plasma cells within colon lamina propria of 10- and 26-week-old mice. **(f-j)** Uninfected *Tbx21^tdTomato-creERT2^ Rosa26^LSL-YFP^* were administered tamoxifen by oral gavage at 5 or 12 weeks of age and analyzed by flow cytometry 6.5 weeks following tamoxifen administration. **(f)** Frequency of T-bet recent (tdTomato+ YFP-), T-bet transient (tdTomato-YFP+), and T-bet stable (tdTomato+ YFP+) cells within reporter+ MBC from mice given a tamoxifen pulse at 5 weeks of age. **(g)** Frequency of T-bet recent (tdTomato+ YFP-), T-bet transient (tdTomato- YFP+), and T-bet stable (tdTomato+ YFP+) cells within reporter+ PC from mice given a tamoxifen pulse at 5 weeks of age **(h)** Representative two-dimensional flow cytometry plots pre-gated on MBC. Flow plots show gating strategy for identifying tdTomato expression within YFP+ B cell subsets. **(i)** Frequency of tdTomato+ cells within YFP+ MBC and **(j)** YFP+ PC in mLN, PP, and lamina propria (small intestine (SI) and colon) from mice given a tamoxifen pulse at 5 and 12 weeks of age. **(d-e & i-j)** Each dot represents a mouse, bars show mean, error bars show S.D; 2-way ANOVA with Tukey’s correction for multiple comparisons. **(f-g)** Bars show mean, error bars show S.D. n=3-4.

**Extended data Figure 2.**
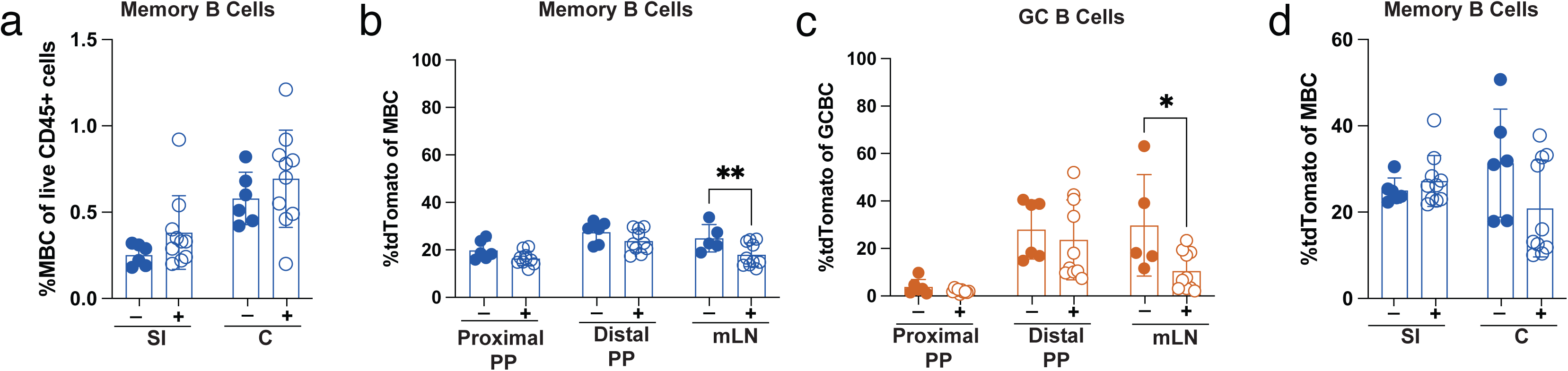
*Tbx21^tdTomato-cre^* mice were administered triple antibiotics in drinking water for 14 days. **(a)** Frequency of memory B cells (MBC) in small intestine (SI) and colon lamina propria. **(b)** Frequency of tdTomato+ memory B cells (MBC) in mLN and PP. **(c)** Frequency of tdTomato+ GC B cells (GCBC) in mLN and PP**.(d)** Frequency of tdTomato+ memory B cells (MBC) in small intestine (SI) and colon lamina propria. **(a-d)** Each dot represents a mouse, bars show mean, error bars show S.D; 2-way ANOVA with Bonferroni correction for multiple comparisons *P< 0.033(*), P< 0.002(**)*.

**Extended data Figure 3.**
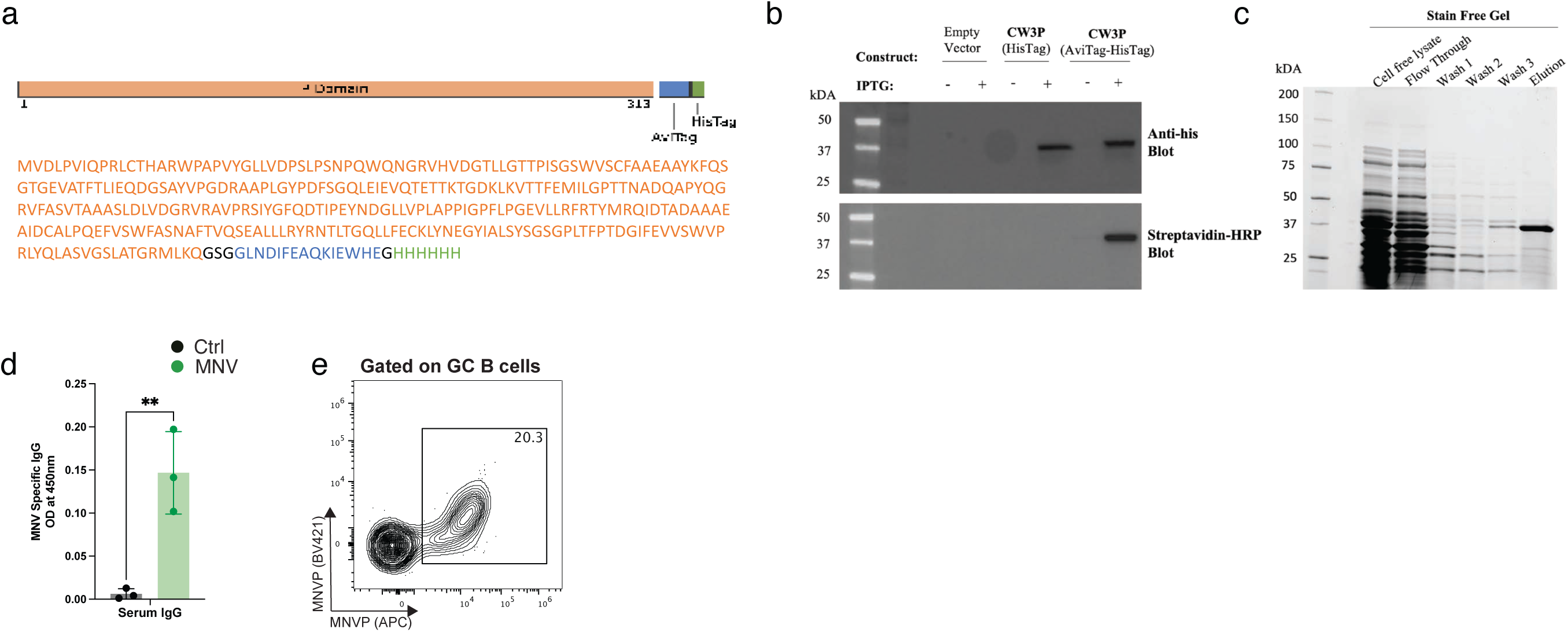
**(a)** Amino acid sequence of the MNVP probe construct. **(b)** Anti-his and Streptavidin HRP western blot showing his-tag incorporation and biotinylation of expressed MNVP probe. **(c)** Stain free gel showing purity of MNVP after Nickel-Nitrilotriacetic Acid resin (NiNTA) affinity purification. **(d)** MNV specific serum IgG titers against MNVP capture 21 dpi. **(e)** Representative two-dimensional flow cytometry plot showing dual probe stain on GC B cells (GCBC), x-axis (MNVP-APC) and y-axis (MNVP-BV421). Each dot represents a mouse, bars show mean, error bars show S.D; Unpaired t-tests. *P< 0.002(**)*.

**Extended data Figure 4.**
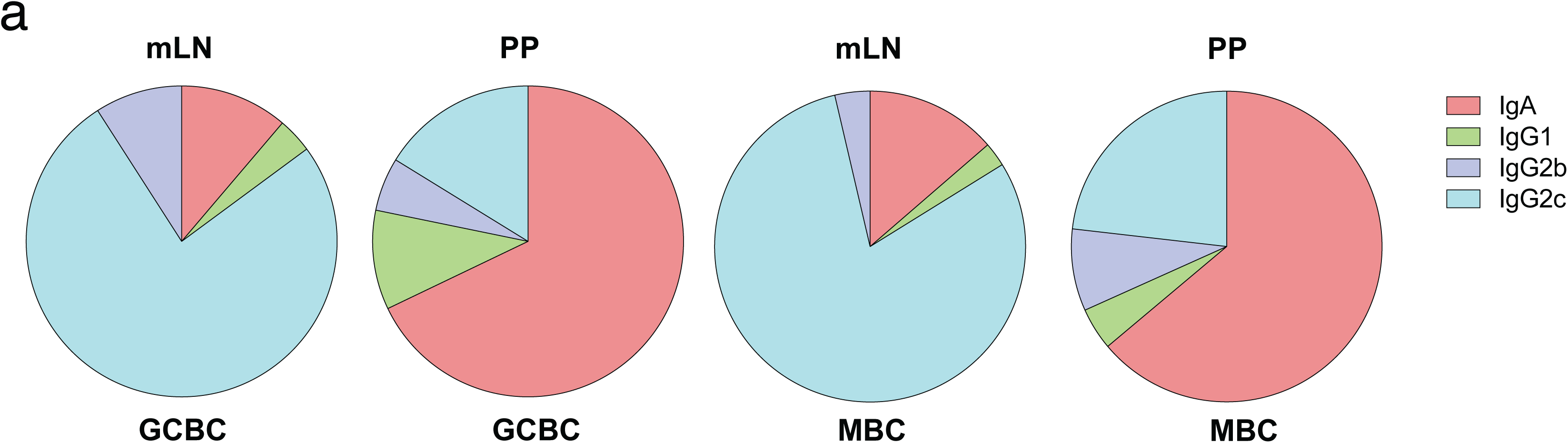
*Tbx21^tdTomato-cre^*mice were infected with 1E6 PFU of MNV and lymphoid organs profiled at 14dpi. **(a)** Frequency of antibody isotypes within MNVP+ GC B cells (GCBC) and memory B cells (MBC).

**Extended data Figure 5.**
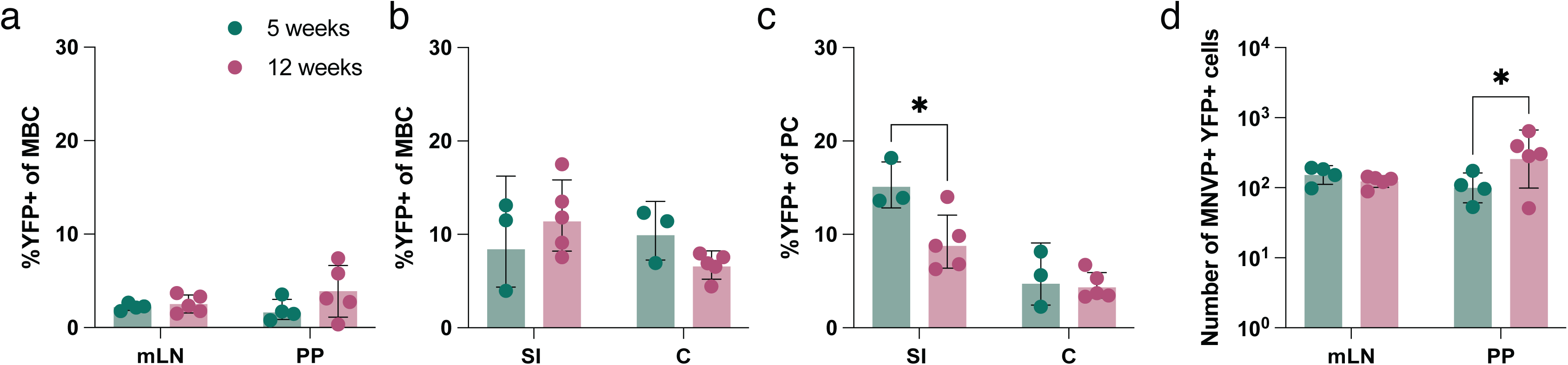
*Tbx21^tdTomato-creERT2^ Rosa26^LSL-YFP^* mice were infected with 1E6 PFU of MNV and administered tamoxifen 14 dpi by oral gavage, indicated organs were analyzed at indicated timepoints by flow cytometry. **(a)** Frequency of YFP+ memory B cells (MBC) in mLN and PP at 5 and 12 weeks post labelling. **(b)** Frequency of YFP+ memory B cells (MBC) in lamina propria of small intestine and colon at 5 and 12 weeks post labelling. **(c)** Frequency of YFP+ plasma cells (PC) in lamina propria of small intestine and colon at 5 and 12 weeks post labelling. **(d)** Number of MNVP+ YFP+ memory B cells (MBC) in mLN and PP 5 and 12 weeks post labelling. (**a-d**) Each dot represents a mouse, bars show mean, error bars show S.D; 2-way ANOVA with Bonferroni correction for multiple comparisons *P< 0.033(*)*.

**Extended data Figure 6.**
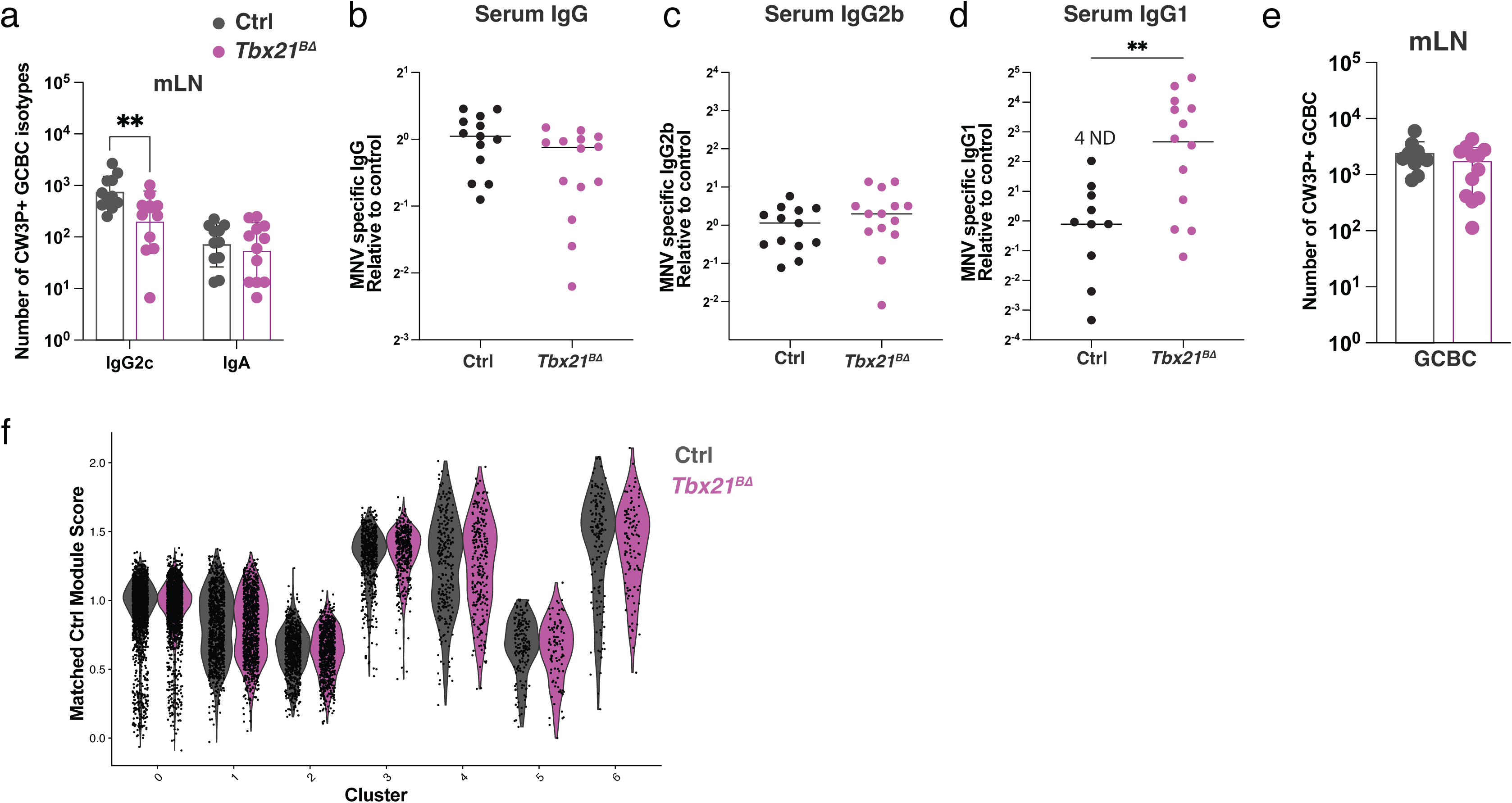
**(a-e)** *Tbx21^fl/fl^ CD23^Cre^* (*Tbx21^BΔ^*) and control *Tbx21^fl/wt^ CD23^Cre^* (Ctrl) mice were infected with 1E6 PFU of MNV and analyzed at 21 dpi by flow cytometry or ELISA. **(a)** Number of MNVP+ IgG2c and IgA switched GC B cells (GCBC) in mLN. **(b-d)** MNV-specific antibodies were quantified by ELISA in serum at 21 dpi **(b)** IgG **(c)** IgG2b **(d)** IgG1. Absorbance was normalized to control and displayed as fold change on log scale. **(e)** Number of MNVP+ GC B cells (GCBC) in mLN. **(f)** Ctrl derived cluster specific module scores applied to both Ctrl and *Tbx21^BΔ^* cells. **(a-e)** Each dot represents a mouse, bars show mean, error bars show S.D; **(a)** 2-way ANOVA with Bonferroni correction for multiple comparisons; **(b-c & e)** Unpaired two-tailed t-test; **(d)** Mann-Whitney test; **(f)** Wilcoxon Rank Sum test. *P< 0.002(**)*.

**Extended data Figure 7.**
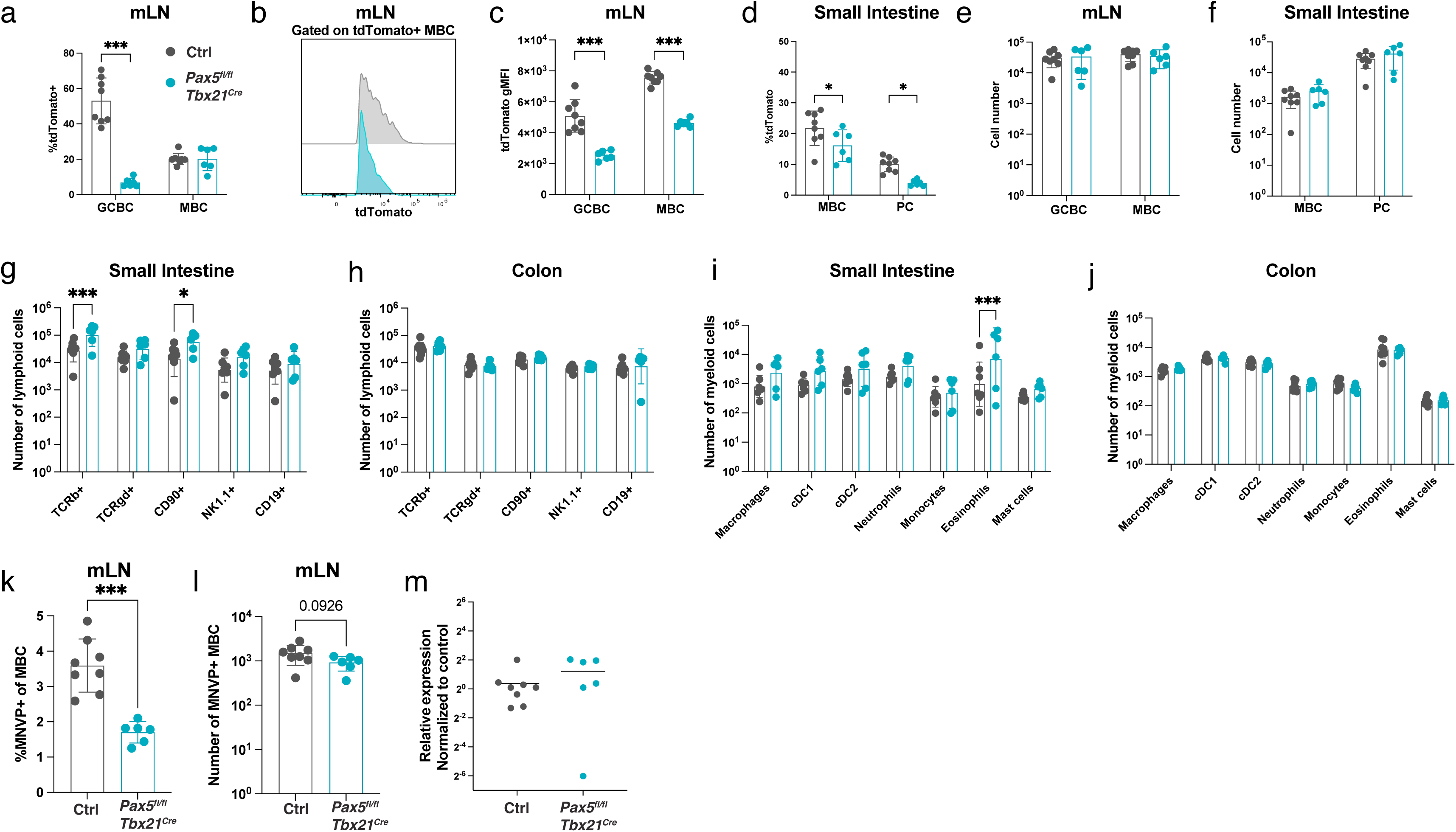
**(a-l)** *Pax5^fl/fl^ Tbx21^Cre^* and control *Pax5^fl/wt^ Tbx21^Cre^* (Ctrl) mice were infected with 1E6 PFU of MNV and organs analyzed 21 dpi or **(m)** at 3 dpi. **(a)** Frequency of tdTomato+ GC B cells (GCBC) and memory B cells in mLN and PP. **(b)** Representative two-dimensional flow-cytometry plot showing fluorescence intensity of tdTomato in tdtomato+ memory B cells (MBC). **(c)** Geometric mean fluorescence intensity of tdTomato expression within tdTomato+ GC B cells (GCBC) and memory B cells (MBC) in mLN. **(d)** Frequency of tdTomato memory B cells (MBC) and plasma cells (PC) in small intestine lamina propria. **(e)** Number of GC B cells (GCBC) and memory B cells (MBC) in mLN. **(f)** Number of memory B cells (MBC) and plasma cells (PC) in small intestine. **(g)** Number of lymphoid cells in small intestine lamina propria. **(h)** Number of lymphoid cells in colon lamina propria. **(i)** Number of myeloid cells in small intestine lamina propria. **(j)** Number of myeloid cells in colon lamina propria. **(k)** Frequency and **(l)** number of MNVP+ memory B cells (MBC) in mLN at 21 dpi. **(m)** Relative MNV RNA abundance in the distal ileum at 3 dpi, determined by RT-qPCR. Data are presented as fold change relative to controls, calculated using the 2^-ΔΔCT^ method with Actb as the internal reference gene. Each dot represents a mouse, bars show mean, error bars show S.D.; **(a, c-j)** 2-way ANOVA with Bonferroni correction for multiple comparisons; **(k-m)** Unpaired two-tailed t-test. *P< 0.033(*), P < .001(***)*.

